# Genome assembly and genomic architecture of a prominent cold-resistant rapeseed germplasm

**DOI:** 10.1101/2023.11.12.566742

**Authors:** Zefeng Wu, Guoqiang Zheng, Yali Sun, Xiaoyun Dong, Ying Wang, Hui Li, Jiaping Wei, Junmei Cui, Yan Fang, Yinin Niu, Zhen Huang, Jihong Hu, Zigang Liu

## Abstract

The genomic landscape of cold-tolerant winter rapeseed (*Brassica napus*, L) has been poorly characterized. We assembled a high-quality reference genome of a prominent cold-tolerant winter rapeseed cultivar, NTS57, and performed phylogenetic and pan-genomic analyses by integrating other reported *Brassica napus* accessions. The transcriptome analysis revealed that microtubule-associated biological pathways were much more active in NTS57 under cold stress than in the cold-sensitive variety. Whole genome methylation data analysis revealed that DNA demethylation on protein-coding genes and repetitive elements, especially at CHH sites, is essential for cold response in winter rapeseed.

---

To the editor,

Winter rapeseed is an important oilseed crop that is often subjected to low temperature stress and consequent yield loss at high latitudes. Therefore, identification of highly cold-resistant winter type rapeseed and elucidation of the genomic architecture are crucial for genetic improvement of the cold-resistant trait. Here, we assembled a chromosome-level reference genome of the prominent cold-resistant winter rapeseed cultivar NTS57 by integrating Illumina, PacBio and Hi-C sequencing data. Phylogenetic and comparative genomic analyses were performed to discover the relationships and genomic variations between NTS57 and other rapeseed accessions. Transcriptomic analysis of NTS57 and a cold-sensitive cultivar under cold stress identified > 10,000 differentially expressed genes that were enriched in regulation of circadian rhythm, photosynthesis, and cold response. Many important molecular components, including *DREBs, ZAT12, CAMTA3/5*, and *CORs*, in the classical cold signaling pathway were also discovered. Importantly, genes or biological pathways associated with microtubules were much more active in NTS57 under cold stress relative to the cold-sensitive cultivar, underlying its prominent cold resistance. We also showed that methylation levels were significantly decreased under cold stress, suggesting that demethylation may be required for the cold stress response. The high-quality NTS57 rapeseed reference genome and other omics data will be valuable genetic resources for understanding cold tolerance in rapeseed.

## RESULTS AND DISCUSSIONS

### Genome assembly and annotation of a cold tolerant rapeseed NTS57

*Brassica napus* NTS57 (2n = 19, AACC) is a cultivated rapeseed developed by crossing winter rapeseed with *B. rapa*. It has strong cold resistance and can survive in the field at −26°C in northwest China. To explore the genetic architecture that contributes to the cold resistance of NTS57, we performed DNA sequencing and de novo assembly of the NTS57 genome by integrating 60 × PacBio high-fidelity (HiFi) long-read sequencing, 100× Hi-C sequencing, and 30 × Illumina paired-end sequencing data. First, K-mer analysis of the Illumina sequencing reads estimated the genome size of NTS57 to be 1.01–1.29 Gb (Figure S1). Then, the PacBio reads were assembled and polished into a 1.06-Gb genome assembly with contig N50 of 21.81 Mb, and > 93.6% (995 Mb) of the sequences were anchored on 19 chromosomes with scaffold N50 of 58.64 Mb after Hi-C scaffolding (Figure 1A, B). Centromere sequences were successfully identified for all 19 NTS57 chromosomes (Figure 1B and Table S1). The high completeness of the genome assembly was evidenced by a BUSCO recovery score of 99.75%. The assembly quality of the present sequenced genome is higher than those of the published rapeseed reference genomes (Table S2). We identified 101,438 protein-coding genes in the NTS57 genome using ab initio prediction, homology search, and transcriptome mapping strategies. Among them, 96,922 (95.5%) were located on 19 chromosomes; 44,411 on the A subgenome and 52,511 on the C subgenome, which is close to the gene numbers in the Darmor*-bzh* and ZS11 genomes (Table S3). By homology search, 97.89% were annotated by at least one of the protein-related databases (Figure S2). In addition, 36,552 non-coding RNAs were annotated in the NTS57 genome (Table S4). Annotation of the NTS57 genome showed that > 55.6% of the genomic content was repetitive elements (Table S5); 80% of them were TEs, including retrotransposons and DNA transposons. The Gypsy and Copia represented the highest proportion of the retrotransposons (Table S5). Genomic visualization showed that Gypsy and Copia had overall similar distribution patterns along the chromosomes and an anti-correlation with gene density; however, they displayed an opposite pattern near centromeres with high GC content and DNA methylation level (Figure 1B), which is similar to the findings of a previous study [1]. Chromosomal synteny analysis based on protein-coding gene arrangements detected strong intraspecific collinearity between homologous chromosomes from the two subgenomes (Figure 1B), demonstrating the homology of chromosomes that were inherited from two ancestral diploid genomes.

**Figure 1.**
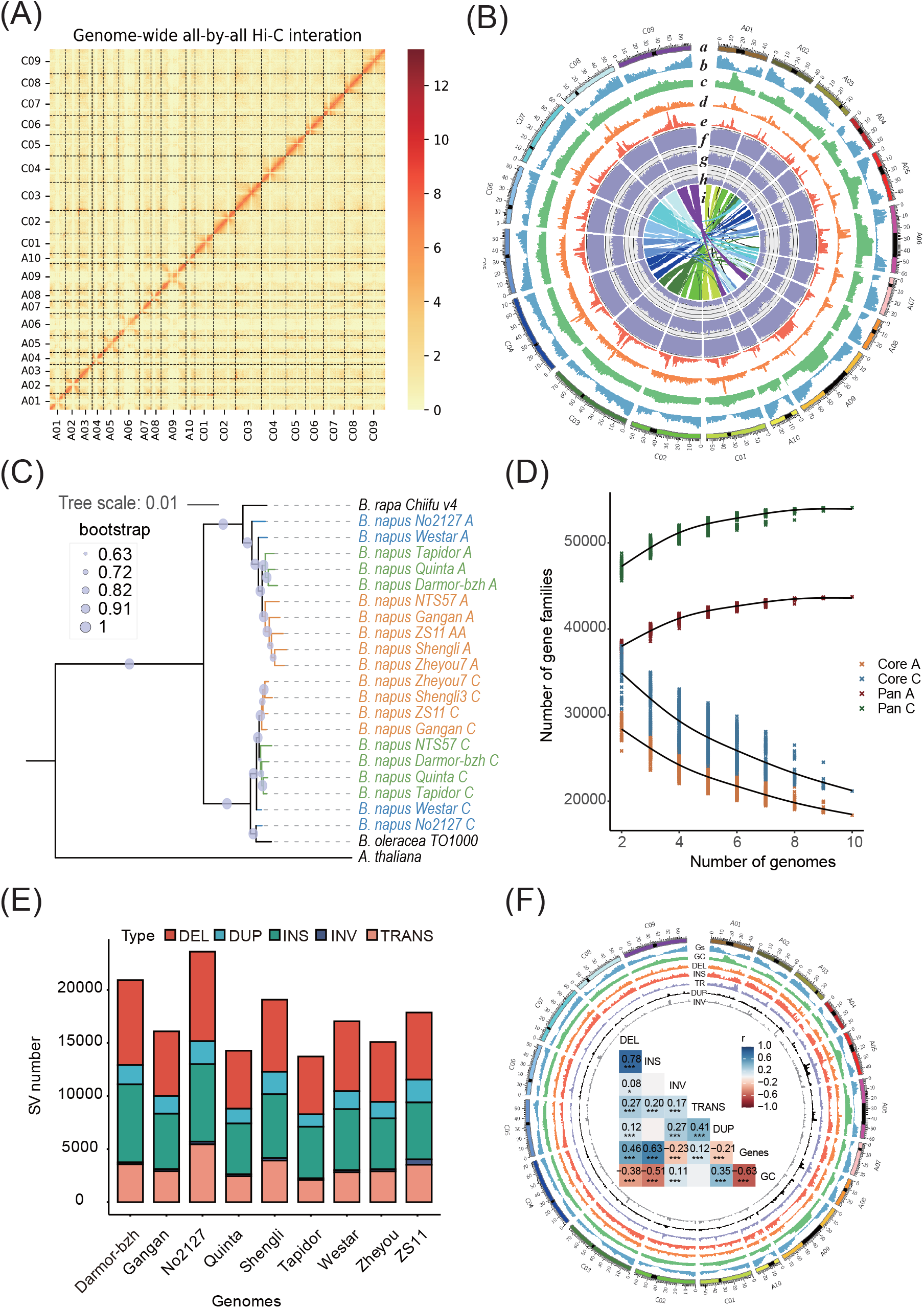
Genome assembly of NTS57 and pangenome analysis of rapeseed. (A) Genome-wide Hi-C heat map of the NTS57 genome. The diagonal pattern indicates strong Hi-C intra-chromosomal interactions of the A and C subgenomes of NTS57. The color bar indicates the log-transformed interaction frequencies of the Hi-C chromosome links. (B) Circos plot showing the genome assembly and genomic features of NTS57. The letters (*a–i*) indicate the ideograms of 19 NTS57 chromosomes, gene density, GC content, Copia transposons, Gypsy transposons, CpG methylation, CHG methylation, CHH methylation, and chromosomal collinearity, respectively. The black rectangles in the ideograms indicate the centromeres. (C) Maximum likelihood tree constructed using 2,268 single-copy orthologous genes shows the phylogenetic relationships of NTS57 and other public rapeseed accessions. Bootstrap values inferred from 1000 replicates are given as percentages on the branches of the consensus tree. (D) Modelling of pan-genome and core genome sizes by including additional rapeseed accessions. (E) Number of different structural variation (SV) types in the genome of each rapeseed accession using NTS57 as the reference genome. DEL, DUP, INS, INV, and TRANS, deletion, duplication, insertion, inversion, and translocation, respectively. (F) SV landscape in the NTS57 genome. The heat map shows the correlations between the different SVs, GC content, and gene density (Gs). The correlation significance is marked in the corresponding cells.

### Phylogeny and pan-genomic analysis of rapeseed varieties

To determine the phylogeny of NTS57 with other rapeseed cultivars, we integrated nine publicly sequenced rapeseed accessions with different ecotypes, namely three winter type (Darmor-*bzh*, Quinta, and Tapidor), four semi-winter type (ZS11, Shengli, Zheyou, and Gangan), and two spring-type (Westar and No2127) rapeseed cultivars. The maximum likelihood tree constructed using 2,268 single-copy orthologous genes showed that the A subgenome of NTS57 was closely related to the semi-winter rapeseed accessions, whereas the C subgenome formed a clade with the winter type rapeseed accessions (Figure 1C). This result indicated that NTS57 and semi-winter rapeseed bred in China had different breeding histories, and the cold tolerance may have been inherited or retained from winter type rapeseed. We then investigated the pan-genome architecture of rapeseed by integrating NTS57 and nine other rapeseed accessions. By clustering protein-coding genes, we identified 43,734 and 54,091 pangene families from the A and C subgenomes, respectively. The number of gene families identified in the A and C subgenomes increased rapidly when more subgenomes or species were incorporated (Figure 1D), implying that the 10 rapeseed genomes have rich gene pools and that a single reference genome cannot capture the full genetic diversity of rapeseed. For the A subgenome, more than 18,375 gene families were conserved among the 10 rapeseed accessions, and were considered to represent the core gene families. Dispensable gene families that were present in 2–10 accessions accounted for 55.69% of the pangene families, and 2.3% of the pangene families were categorized as genome specific (Figure S3A). For the C subgenome, similar percentages of core, dispensable, and genome-specific pangene families were identified, although there was a slightly lower percentage of core gene families and a slightly higher percentage of dispensable gene families (Figure S3B). Gene function enrichment analysis showed that the core gene families and the indispensable gene families have different functions in rapeseed growth and development (Figure S4).

The high-quality NTS57 reference genome allowed the SVs associated with cold tolerance to be evaluated by comparative genomic analysis. A total of 51,511 insertions, 58,775 deletions, 29,645 translocations, 15,654 duplications, and 4,047 inversions were identified in the nine rapeseed accessions with NTS57 as the reference genome (Figure 1E). The artificially synthesized rapeseed line No2127 had the highest number of SVs, and Tapidor had the smallest number of SVs. The total sizes of the SVs were 113.5–153.6 Mb in the different accessions, with the length per SV ranging from a few tens of bp to hundreds of kb or even above the Mb scale (Figure S5). We also observed that insertions and deletions tended to be enriched at both chromosome ends and depleted near the centromeres, with a distribution similar to that of gene density and opposite to that for GC content. Unlike the deletions and insertions, the inversions, duplications, and translocations were randomly distributed across the 19 chromosomes, although some enrichment was detected near centromeres (Figure 1F and Figure S6). These findings were confirmed by calculating the correlations between the chromosomal distributions of the different SVs (Figure 1F).

### Gene dynamic expression and DNA methylation of NTS57 under cold stress

To identify genes associated with cold tolerance in NTS57, we analyzed the transcriptomic data of plants under normal and cold stress conditions that had been generated in a previous study [2]. Compared with the control, we identified 15,888 and 22,479 DEGs in the T1 (12h after cold) and T2 (24h after cold) samples, respectively (Figure 2A and Table S6, S7). As expected, the number of identified cold-responsive genes was higher at the longer time after cold stress. We identified 12,346 DEGs that were common at both time points after cold stress (Figure S7), suggesting that these genes may be core molecular components in NTS57 in response to cold stress. We found that many DEGs were directly involved in plant cold response or cold acclimation processes, including genes that encoded superoxide dismutase (SOD) [3], VERNALIZATION INSENSITIVE 3 (VIN3) [4], transcription factor LUX [5], RNA-binding protein CP29B [6], flavanone 3-dioxygenase 3 [7], and dehydration-responsive element-binding protein 2E (DREB2E) [8]. Furthermore, many DEGs were annotated as cold-responsive or cold-induced genes (Table S6,S7). We then performed GO and KEGG pathway enrichment analysis. The DEGs in T1 were annotated mainly with regulation of circadian rhythms, cold response, and xyloglucan metabolism terms under the biological process category, MCM complex, cell wall, and microtubule under the cellular component category, and UDP-glycosyltransferase, protein serine/threonine phosphatase activity, and ubiquitin protein ligase activity under the molecular function category (Figure 2B). The DEGs in T2 were enriched with the same GO terms as the DEGs in T1, as well as photosynthesis and monopolar cell growth under the biological process category (Figure S8), suggesting that more biological processes are affected in the response to cold stress with longer stress duration. The KEGG pathway enrichment results showed that the DEGs were involved mainly in MAPK signaling pathway, circadian rhythm, flavonoid biosynthesis, and photosynthesis, which is similar to the results of the GO enrichment analysis (Figure S9). To identify potential genes and pathways specific to the prominent cold tolerance rapeseed NTS57, we performed similar gene expression analysis on the cold-sensitive rapeseed NF cultivar, which was found to have an extremely low overwintering survival rate based on previous phenotype observation and physiological measurements [2]. Principal component analysis showed that NTS57 and NF samples under cold stress were separated from each other, suggesting that accession-specific transcriptomic changes occurred upon cold treatment (Figure S10). The comparative transcriptomic analysis at the whole genome level showed that there were high correlations in gene expression changes between the NTS57 and NF accessions in both T1 (*R* = 0.68) and T2 (*R* = 0.79) samples (Figure S11). We identified 15,558 and 20,988 high-quality DEGs in the T1 and T2 samples of NF, respectively (Table S8, S9), which is comparable with that in the NTS57. Besides, many DEGs in the NF were overlapped with those identified in the NTS57 (Figure S12). Similar gene function enrichments were also observed for the DEGs of NF derived from the two stress time points (Figure 2B and Figures S8, S13), implying that these biological processes are essential or conserved across rapeseed accessions under cold stress. Despite the high similarity of transcriptomic changes between the two accessions, a number of genes were exclusively expressed in NTS57 but not in NF in T1 or T2 samples (Figure S11), which may explain the difference in cold tolerance between the two accessions. The enrichment analysis results showed that genes involved in enriched microtubule-associated processes under the biological process, cellular component, and molecular function GO categories were all specifically altered in NTS57 upon cold stress (Figure S14). Gene expression profiles of microtubule-associated genes also changed more dramatically in NTS57 than they did in NF under cold stress, especially *A05G015370* (*TUBB7*), *A06G029560* (*TOR1L3*), and *C05G002130* (*TPX2*) (Figure 2C). These finding indicate that microtubule-associated processes were more active in NTS57 than they were in NF, and likely contributed to the cold tolerance in NTS57.

**Figure 2.**
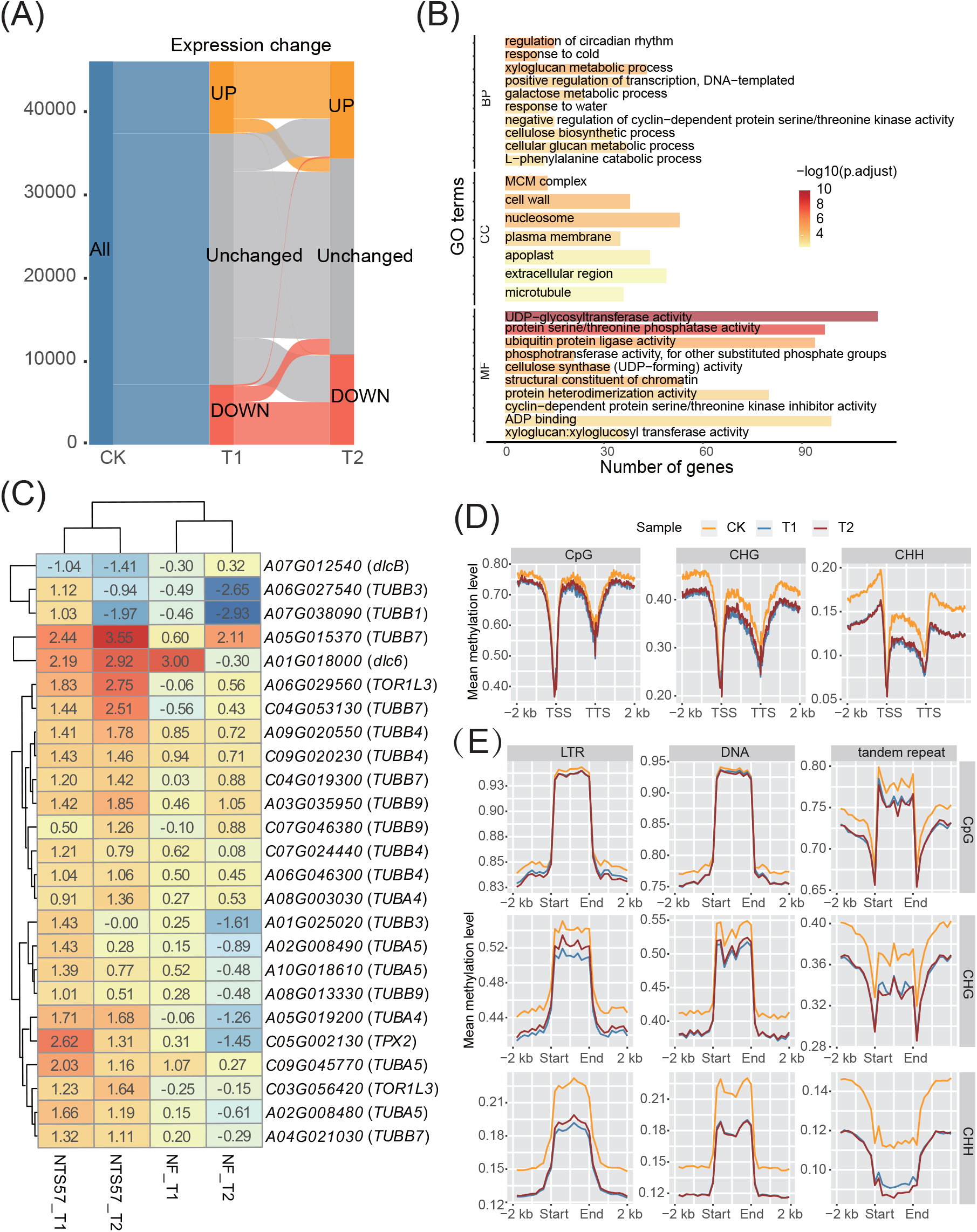
Gene dynamic expression and DNA methylation of NTS57 under cold stress. (A) Sankey plot showing the up- and down-regulated genes of NTS57 after 12 h (T1) and 24 h (T2) cold stress at −4°C. CK, normal conditions at 25°C. (B) Gene ontology (GO) function enrichment of DEGs in the T1 samples. BP, biological process; CC, cellular component; MF, molecular function. (C) Heat map of the expression change of the NTS57-specific DEGs involved in microtubule process in accessions NTS57 and NF under cold stress. (D) Profiles of methylation levels in the gene body, and upstream and downstream regions in the NTS57 genome under normal and cold stress. CK, normal condition at 25°C; T1 and T2, cold stress at −4°C for 12 h and 24 h, respectively; TSS, transcription start site; TTS, transcription termination site; LTR, long terminal repeat. The gene body was divided into 100 bins of equal size. For each bin in each gene, the methylation level was calculated by dividing the total number of methylated Cs by the total number of Cs, and the average of methylation level in the window across all protein-coding genes was used to generate the profiles. The three methylation profiles for the CpG, CHG, and CHH contexts were analyzed independently. (E) Profiles of methylation levels in the repetitive elements of NTS57. CK, T1, and T2 are as defined in (D).

Cold stress has been reported to induce genomic demethylation in many plant species [9]. By calculating the genome-wide DNA methylation levels of the NTS57 genome, we found a remarkable decrease in methylation levels across the gene body and flanking regions after cold stress in all cytosine sequence contexts, namely CpG, CHG, and CHH (Figure 2D). In particular, the decrease in CHH methylation was most pronounced, followed by CHG and CpG methylation, implying that CHH methylation was more sensitive to cold stress. When the A and C subgenomes were treated independently, similar trends were observed (Figure S15), which is consistent with the findings of previous studies [10, 11]. In plants, DNA methylation is enriched over heterochromatic TEs and repetitive elements [12]. As expected, there was high enrichment of DNA methylation on repeat sequences, including long terminal repeats, DNA transposons, and tandem repeats in the NTS57 genome (Figure 2E). However, we noticed two exceptions for CHG and CHH methylation in tandem repeats, which showed some deletions compared with the surrounding regions (Figure 2E). We also found that methylation levels were decreased in all the repeat sequences and in all methylation contexts in NTS57 under cold stress, which is very similar to that of the genic regions, suggesting that methylase acts on genes and repeats in a similar way in the plant cold stress response.

## AUTHOR CONTRIBUTIONS

**Zefeng Wu**: Conceptualization, Funding acquisition, Formal analysis, Methodology, Visualization, Writing - original draft. **Guoqiang Zheng**: Conceptualization, Formal analysis, Investigation. **Yali Sun:** Conceptualization, Formal analysis. **Xiaoyun Dong:** Visualization, Software, Validation. **Ying Wang:** Software, Investigation. **Hui Li:** Investigation, Validation. **Jiaping Wei:** Investigation, Validation. **Junmei Cui:** Visualization, Software. **Yan Fang:** Investigation, Validation. **Yinin Niu:** Investigation, Validation. **Zhen Huang:** Investigation, Writing – review & Editing. **Jihong Hu:** Conceptualization, Writing – review & Editing. **Zigang Liu:** Conceptualization, Funding acquisition, Project administration, Supervision.

## ACKNOWLEDGMENTS

This work was supported by the Research Program Sponsored by State Key Laboratory of Aridland Crop Science, Gansu Agricultural University (GSCS-2023-03), the Scientific Research Starting Foundation of Gansu Agricultural University (GAU-KYQD-2020-27), the Natural Science Foundation of Gansu Province (22JR5RA862 and 23JRRA1421), the Gansu Province Higher Education Institutions Young Doctoral Fund (2023QB-126), the Gansu Provincial Science and Technology Major Project (22ZD6NA009), the Central-Guided Local Science and Technology Development Fund (ZCYD-2021-17), and the National Natural Science Foundation of China (32360520).

## DATA AVAILABILITY STATEMENT

All the raw sequencing data and the genome assembly generated for this research are available in the NCBI BioProject database under accession number PRJNA1012021. Genome annotation files and other related documents are available in figshare (https://doi.org/10.6084/m9.figshare.24098538.v1) or in the public database (http://oilseed.online:8000/). The data and scripts used are saved in GitHub https://github.com/Zefeng2018/NTS57_genome_assembly. Supplementary materials (methods, figures, tables, graphical abstract, slides, videos, Chinese translated version and update materials) may be found in the online DOI or iMeta Science http://www.imeta.science/imetaomics/.

## ETHICS STATEMENT

No animals or humans were involved in this study.

## SUPPORTING INFORMATION

The online version contains supplementary figures and tables available.

Figure S1. Genome size estimation based on survey sequencing.

Figure S2. Venn diagram of gene function annotations obtained using several protein-related databases.

Figure S3. Percentage of core genes, dispensable genes, and accession-specific genes in the A and C subgenomes of rapeseed.

Figure S4. Gene function enrichment for core and dispensable genes of NTS57.

Figure S5. Length statistics for structural variations (SVs) in different rapeseed accessions using NTS57 as the reference genome.

Figure S6. Synteny and rearrangement plot for 19 chromosomes of NTS57 and ZS11. Figure S7. Number of DEGs identified in NTS57 under cold stress.

Figure S8. Gene ontology (GO) functional enrichment for the DEGs identified in T2 samples relative to those in the normal condition samples.

Figure S9. KEGG pathway enrichment for the DEGs identified in T1 or T2 samples relative to those in normal condition (CK) samples.

Figure S10. Principal component analysis of gene expression profiles from accessions NTS57 and NF under cold stress.

Figure S11. Gene expression changes in NTS57 and NF under cold stress.

Figure S12. Veen diagram showing the intersection of the DEGs in NTS57 and NF in the T1 and T2 samples.

Figure S13. Gene functional enrichment of the DEGs in NF in the T1 and T2 samples.

Figure S14. Gene functional enrichment of the NTS57-specific differentially expressed genes (DEGs) of NTS57 in the T1and T2 samples.

Figure S15. Methylation level profiles of the A and C subgenomes of NTS57 under cold stress (T1 and T2) and normal conditions (CK).

Table S1. Distribution of predicted centromeres in the NTS57 reference genome.

Table S2. Summary of genome assembly and annotation of NTS57 and other public rapeseed genomes..

Table S3. Comparison of gene number between different rapeseed accessions.

Table S4. Summary of non-coding RNAs in the NTS57 genome.

Table S5. Summary of repetitive elements in the NTS57 genome.

Table S6. Summary of differentially expressed genes in the NTS57 after 12 hours of cold exposure.

Table S7. Summary of differentially expressed genes in the NTS57 after 24 hours of cold exposure.

Table S8. Summary of differentially expressed genes in the NF after 12 hours of cold exposure.

Table S9. Summary of differentially expressed genes in the NF after 24 hours of cold exposure.

## REFERENCES

1. Chen, Xuequn, Chaobo Tong, Xingtan Zhang, Aixia Song, Ming Hu, Wei Dong, Fei Chen, et al. 2021. “A high-quality Brassica napus genome reveals expansion of transposable elements, subgenome evolution and disease resistance.” Plant Biotechnol J 19: 615–630. 10.1111/pbi.13493

2. Wei, Jiaping, Guoqiang Zheng, Xingwang Yu, Sushuang Liu, Xiaoyun Dong, Xiaodong Cao, Xinling Fang, et al. 2021. “Comparative Transcriptomics and Proteomics Analyses of Leaves Reveals a Freezing Stress-Responsive Molecular Network in Winter Rapeseed (Brassica rapa L.).” Front Plant Sci 12: 664311. 10.3389/fpls.2021.664311

3. Lin, Kuan-Hung, Sin-Ci Sei, Yu-Huei Su, Chih-Ming Chiang. 2019. “Overexpression of the Arabidopsis and winter squash superoxide dismutase genes enhances chilling tolerance via ABA-sensitive transcriptional regulation in transgenic Arabidopsis.” Plant Signal Behav 14: 1685728. 10.1080/15592324.2019.1685728

4. Zhao, Yusheng, Rea L Antoniou-Kourounioti, Grant Calder, Caroline Dean, Martin Howard. 2020. “Temperature-dependent growth contributes to long-term cold sensing.” Nature 583: 825–829. 10.1038/s41586-020-2485-4

5. Chow, Brenda Y, Sabrina E Sanchez, Ghislain Breton, Jose L Pruneda-Paz, Naden T Krogan, Steve A Kay. 2014. “Transcriptional regulation of LUX by CBF1 mediates cold input to the circadian clock in Arabidopsis.” Curr Biol 24: 1518–1524. 10.1016/j.cub.2014.05.029

6. Kupsch, Christiane, Hannes Ruwe, Sandra Gusewski, Michael Tillich, Ian Small, Christian Schmitz-Linneweber. 2012. “Arabidopsis chloroplast RNA binding proteins CP31A and CP29A associate with large transcript pools and confer cold stress tolerance by influencing multiple chloroplast RNA processing steps.” Plant Cell 24: 4266–4280. 10.1105/tpc.112.103002

7. Wu, Lishuang, Jian Tian, Yao Yu, Lixia Yuan, Yujiong Zhang, Hao Wu, Furong Wang, Xin Peng. 2023. “Functional characterization of a cold related flavanone 3-hydroxylase from Tetrastigma hemsleyanum: an in vitro, in silico and in vivo study.” Biotechnol Lett 10.1007/s10529-023-03440-5

8. Agarwal, Pradeep K, Kapil Gupta, Sergiy Lopato, Parinita Agarwal. 2017. “Dehydration responsive element binding transcription factors and their applications for the engineering of stress tolerance.” J Exp Bot 68: 2135–2148. 10.1093/jxb/erx118

9. Peng, Hai, Jing Zhang. 2009. “Plant genomic DNA methylation in response to stresses: Potential applications and challenges in plant breeding.” Progress in Natural Science 19: 1037–1045. 10.1016/j.pnsc.2008.10.014

10. Bird, Kevin A, Chad E Niederhuth, Shujun Ou, Malia Gehan, J Chris Pires, Zhiyong Xiong, Robert VanBuren, Patrick P Edger. 2021. “Replaying the evolutionary tape to investigate subgenome dominance in allopolyploid Brassica napus.” New Phytol 230: 354–371. 10.1111/nph.17137

11. Chalhoub, Boulos, France Denoeud, Shengyi Liu, Isobel A P Parkin, Haibao Tang, Xiyin Wang, Julien Chiquet, et al. 2014. “Early allopolyploid evolution in the post-Neolithic Brassica napus oilseed genome.” Science 345: 950–953. 10.1126/science.125343

12. Gallego-Bartolome, Javier. 2020. “DNA methylation in plants: mechanisms and tools for targeted manipulation.” New Phytol 227: 38–44. 10.1111/nph.16529

