## Supplementary figures for "Genome assembly and genomic architecture of a prominent cold-resistant rapeseed germplasm"

**Runing title**：Genome assembly of a cold-resistant rapeseed germplasm

Zefeng Wu^1^, Guoqiang Zheng^1^, Yali Sun^1^, Xiaoyun Dong^1^, Ying Wang^1^, Hui Li^1^, Jiaping Wei^1^, Junmei Cui^1^, Yan Fang^1^, Yinin Niu^1^, Zhen Huang^2^, Jihong Hu^2^, Zigang Liu^1*^

^1^ State Key Laboratory of Aridland Crop Science, Gansu Agricultural University, Lanzhou, 730070, Gansu, China

^2^ State Key Laboratory of Crop Stress Biology for Arid Areas, College of Agronomy, Northwest A&F University, Yangling, 712100, Shaanxi, China.

### Supplemental methods

#### Plant material and DNA extraction

The seeds of rapeseed NTS57 were successfully germinated, and plants were grown in the greenhouse of Gansu Agricultural University at 20°C/18°C and 16-h/8-h day growth conditions. Leaf tissue was collected from a 4-week-old plant, flash frozen, and stored at −70°C. High molecular weight genomic DNA was extracted using the cetyltrimethylammonium bromide (CTAB) method, followed by purification using a GrandOmics genomic kit for regular sequencing according to the manufacturer’s standard operating procedure. DNA degradation and contamination of the extracted DNA was checked on 0.75% agarose gels. DNA purity was determined using a NanoDrop™ One UV-Vis spectrophotometer (Thermo Fisher Scientific, USA); OD260/280 was 1.8–2.0 and OD 260/230 was 2.0–2.2. Finally, the DNA concentration was measured using a Qubit^®^ 3.0 fluorometer (Invitrogen, USA).

#### PacBio HiFi library preparation and DNA sequencing

For long-read sequencing, the extracted DNA was sequenced on a PacBio Sequel II platform (Pacific Biosciences, Menlo Park, CA, USA) using the circular consensus sequencing (CCS) model from two SMRT cells using 15-kb preparation solutions. The main steps for library preparation were: genomic DNA shearing, DNA damage repair, end repair, and A-tailing, ligation with hairpin adapters from a SMRTbell Express Template Prep Kit 2.0 (Pacific Biosciences), nuclease treatment of the SMRTbell library with a SMRTbell Enzyme Cleanup Kit, size selection, and binding to polymerase.

Briefly, an 8 µg DNA per sample was used for the DNA library preparation. The genomic DNA sample was first sheared with g-TUBEs (Covaris, USA) according to the expected size of the fragments for the library. Single strand overhangs were removed, and DNA fragments were damage repaired, end repaired, A-tailed, and ligated with the hairpin adaptor for PacBio sequencing. The library was treated by nuclease with a SMRTbell Enzyme Cleanup Kit and purified with AMPure PB Beads. Target fragments were screened using PippinHT (Sage Science, USA). The SMRTbell library was purified with AMPure PB Beads, and an Agilent 2100 Bioanalyzer (Agilent Technologies, USA) was used to determine the size of the library fragments. Finally, sequencing was performed on a PacBio Sequel II instrument using Sequencing Primer V5 and a Sequel II Binding Kit 2.2 at the GrandOmics Genome Center (Wuhan, China). After sequencing, the resulting BAM files were submitted to the pbccs program (https://ccs.how/) with default parameters. A total of 65.16 Gb HiFi pass reads with average length 15.6 kb were obtained.

#### Preparation of short reads libraries and DNA sequencing

Short-read libraries were prepared using an MGIEasy Universal DNA Library Prep Kit V1.0 (CAT#1000005250, MGI) according to the standard protocol. Briefly, 1 µg of genomic DNA was randomly fragmented by Covaris. The fragmented DNA was selected to an average size of 200–400 bp using MGIEasy DNA Clean Beads (CAT#1000005279, MGI). The selected fragments were end-repaired, 3′ adenylated, and ligated with adapters. The DNA samples were amplified by PCR and the products were purified using MGIEasy DNA Clean Beads (CAT#1000005279, MGI). The double-stranded PCR products were heat denatured and circularized using the splint oligo sequence in the MGIEasy Circularization Module (CAT#1000005260, MGI). The single-strand circle DNA was formatted as the final library and qualified by QC. The qualified libraries were sequenced on a DNBSEQ-T7RS platform (MGI).

#### Transcriptome library preparation and sequencing

Seven different tissues (flower bud, flower, leaf, pod, root, stem, and seed) of NTS57 at different developmental stages (seedling, squaring, flowering, and maturing) were sampled to perform RNA sequencing (RNA-Seq) with three biological replicates (Table S9). Briefly, total RNA was extracted for each sample by grinding the tissue in TRIzol reagent (TIANGEN) on dry ice, then processed according to the manufacturer’s protocol. Poly-A RNAs were enriched from total RNA using a Dynabeads mRNA Purification Kit (Cat. No. 61006, Invitrogen) and fragmented into small pieces using fragmentation reagent in an MGIEasy RNA Library Prep Kit V3.1 (Cat# 1000005276, MGI). First-strand cDNA was synthesized using random primers and reverse transcriptase, followed by second-strand cDNA synthesis. The synthesized cDNA was end-repaired, A-tailed, and ligated to the sequencing adapters according to the library construction protocol. The cDNA fragments were amplified by PCR and purified using MGIEasy DNA Clean Beads (CAT#1000005279, MGI). The library was analyzed on an Agilent Technologies 2100 Bioanalyzer. The double-stranded PCR products were heat denatured and circularized using the splint oligo sequence in the MGIEasy Circularization Module (CAT#1000005260, MGI). The single-strand circle DNA was formatted as the final library. The qualified libraries were sequenced on a DNBSEQ-T7RS platform.

#### Hi-C (high-throughput chromosome conformation capture) library preparation and sequencing

Freshly harvested leaves from 4-week-old plants were cut into 2-cm pieces and vacuum infiltrated in nuclear isolation buffer supplemented with 2% formaldehyde. Cross-linking was stopped by the addition of glycine and additional vacuum infiltration. The fixed tissue was then ground to powder and resuspended in nuclear isolation buffer to obtain a nuclear suspension. The purified nuclei were digested with 100 units of *Dpn*II and labeled by incubation with biotin-14-dCTP. Biotin-14-dCTP was removed from unligated DNA ends due to the exonuclease activity of T4 DNA polymerase. The ligated DNA was sheared into 300–600-bp fragments, then blunt-end repaired and A-tailed, followed by purification by biotin–streptavidin-mediated pull-down. Finally, the Hi-C libraries were quantified and sequenced using an MGI-2000 platform.

#### Genome size and heterozygosity estimation

To understand the genomic characteristics of NTS57, a k-mer analysis was performed using Illumina DNA data prior to genome assembly to estimate the genome size and heterozygosity. Briefly, quality-filtered reads were subjected to 25-mer frequency distribution analysis using the KMC program [1]. By analyzing the 25-mer depth distribution from the clean reads using FindGSE [2] and GenomeScope2.0 [3], we estimated the genome size as G = K-num/K-depth, where K-num is the total number of 25-mers, K-depth is the k-mer depth, and G is the genome size. Finally, the heterozygosity and repeat content of the NTS57 genome were estimated by combining the simulation data results of Arabidopsis with different heterozygosity and the frequency peak distribution of 25-mers.

#### Genome de novo assembly and completeness assessment

The high quality HiFi reads were used to assemble the initial contigs using the Hifiasm (v0.13-r308) package with default parameters [4]. To improve the accuracy of the assembly, the assembled contigs were polished with Nextpolish [5] using short reads with default parameters. Finally, redundancies were removed using Haplomerger2 [6] to generate the primary contigs.

Hi-C reads were used to support the assembly of the genome at the chromosome level. For this, low-quality Hi-C reads (quality scores < 20), adaptor sequences, and sequences <30 bp long were filtered out using fastp [7], then the clean paired-end reads were mapped to the assembled contig sequences using Bowtie2 (v2.3.2) [8] (-end-to-end --very-sensitive -L 30) to obtain uniquely mapped paired-end reads. Valid interaction paired reads were identified from the unique mapped paired-end reads using [HiC-Pro (v2.8.1)](https://github.com/nservant/HiC-Pro) [9] and retained for further analysis. Invalid read pairs, including dangling-end, self-cycle, re-ligation, and dumped products were filtered. Scaffolds were clustered, ordered, and oriented on chromosomes using LACHESIS [10] with parameters CLUSTER_MIN_RE_SITES=100, CLUSTER_MAX_LINK_DENSITY=2.5, CLUSTER NONINFORMATIVE RATIO = 1.4, ORDER MIN N RES IN TRUNK = 60, ORDER MIN N RES IN SHREDS = 60. Finally, placement and orientation errors with obvious discrete chromatin interaction patterns were adjusted manually. The completeness of the genome assembly was evaluated using BUSCO v4.0.5 (Benchmarking Universal Single-Copy Orthologs) [11]. To assess the accuracy of the assembly, all the Illumina paired-end reads were mapped to the assembled genome using BWA software [12], and the mapping rate and genome coverage of the sequencing reads were assessed using SAMtools v0.1.1855 [13]. The base accuracy of the assembly was calculated using BCFtools [14]. The coverage of potentially expressed genes of the assembly was evaluated by aligning the RNA-Seq data to the assembly using HISAT2 with default parameters [15]. To avoid including mitochondrial sequences in the assembly, the draft genome assembly was queried against the NCBI NT library and sequences that aligned with mitochondrial sequences were eliminated.

#### Repetitive DNA elements and gene model annotation

Tandem repeats were annotated using the GMATA [16] and Tandem Repeats Finder (TRF) [17] software; GMATA can identify simple sequence repeats and TRF can recognize all tandem repeat elements in an entire genome. Transposable elements (TEs) were identified in the NTS57 genome using both de novo and homology-based strategies. For de novo prediction, three de novo prediction programs, MITE-hunter [18], RepeatModeler [19], and LTR_retriever [20], were used with default parameters. The predicted TEs were integrated into a de novo database, then aligned to Repbase (http://www.girinst.org/repbase) and classified into different types. Using the resulting database, repetitive sequences were identified by a homologous sequence search using RepeatMasker with default parameters [21]. Overlapping repeat elements that belonged to the same repeat class were merged to obtain a final non-redundant repeat sequence annotation. To identify centromere sequences among the NTS57 genome sequences, the quarTeT program was used with default parameters [22]. The top candidates were selected based on the length and periods of repetitive sequences. The gene density and long terminal repeat density distributions across the chromosomes were used to help determine the centromere positions in chromosomes.

For gene model prediction, ab initio prediction, homology search, and reference-guided transcriptome assembly were used to predict genes in the repeat-masked genome. GeMoMa [23] was used to align the homologous peptides from related species to the assembly, and gene structure information was obtained by homology prediction. For RNA-Seq-based gene prediction, the filtered RNA-Seq reads were mapped to the reference genome using STAR with default parameters [24]. Then, the transcripts were assembled using StringTie [25], and open reading frames (ORFs) were predicted using PASA (https://github.com/PASApipeline/PASApipeline). For the de novo prediction, RNA-Seq reads were de novo assembled using StringTie [25] and analyzed using PASA to obtain a training set. AUGUSTUS was used with default parameters [26] for ab initio gene prediction with the training set. Finally, EVidenceModeler [27] was used to generate an integrated gene set. Genes with TEs were removed from the integrated set using the TransposonPSI package (http://transposonpsi.sourceforge.net/) and miscoded genes were further filtered. Untranslated regions (UTRs) and alternative splicing regions were determined using PASA based on the RNA-seq assemblies. The longest transcripts for each locus were retained, and regions outside the ORFs were designated as UTRs.

#### Functional annotation of gene models

Functional information of genes, motifs, and domains of their proteins was obtained by searches against public databases, namely Swiss-Prot [28], NCBI nonredundant protein database NR [29], KEGG (Kyoto Encyclopedia of Genes and Genomes) [30], KOG (EuKaryotic Orthologous Groups) [31], and Gene Ontology (GO) [32]. Putative domains and GO terms of the genes were identified using InterProScan with default parameters [33]. BLASTp [34] was used to compare the EvidenceModeler-integrated protein sequences against the Swiss-Prot, NR, KOG, and KEGG databases with an E-value cutoff of 1e−05; the hit with the lowest E-value was retained. All the search results were concatenated.

#### Annotation of non-coding RNAs

Database searches and model prediction were used to identify non-coding RNAs. Transfer RNAs (tRNAs) were predicted using tRNAscan-SE with eukaryote parameters [35]. MicroRNAs, ribosomal RNAs (rRNAs), small nuclear RNAs (snRNAs), and small nucleolar RNAs were detected using the Infernal cmscan program to search the Rfam database [36]. The rRNAs and their subunits were predicted using RNAmmer [37].

#### Synteny analysis between NTS57 and other rapeseed accessions

MCscan (Python version) [38] was used to identify collinear blocks of ≥ 30 genes in the genomes of NTS57 and other rapeseed cultivars.

#### Phylogenetic analysis

To determine the phylogenetic relationship between NTS57 and other rapeseed cultivars, a phylogenetic tree was reconstructed using 10 rapeseed cultivars, with two ancestral reference genomes, *B. rapa* Chiifu (v4.0) and *B. oleracea*, and *Arabidopsis thaliana* as the outgroup species. Single-copy orthogroups of the 13 species were identified using their amino acid sequences by OrthoFinder v2.0.9 with the parameter “-M msa” [39]. Then, multiple sequence alignment of each orthogroup was performed using mafft v7.453 [40]. The resulting alignments were concatenated into a superalignment. The phylogenetic tree was constructed using the maximum likelihood method implemented in IQ-TREE v1.6.9 with 1000 bootstrap tests [41]. The best-fit model was determined as being JTT+F+R4 by ModelFinder with free rate heterogeneity [42].

#### Gene expression analysis

The newly generated RNA-seq datasets and published RNA-seq datasets were used for the gene expression analysis. Trimmomatic (v0.36) [43] with default parameters was used to remove adapters and low-quality reads from the raw RNA-seq data. The clean RNA-seq reads were mapped to the corresponding reference genomes using HISAT2 with default settings [15]. The gene expression level for each transcript was estimated using FPKM (fragments per kilobase of exon per million fragments) values calculated by StringTie (v2.1.6) [25]. For the previous published data [44], the differentially expressed genes (DEGs) between cold stress and control samples were identified using edgeR and DESeq2, with |log_2_(fold change)| ≥1 and false discovery rate < 0.01 [45]. The common DEGs identified by the two programs were used as the final DEGs for further analysis.

#### Analysis of whole-genome bisulfite sequencing data

Whole genome methylation data for leaf tissues of NTS57 under normal conditions at 25°C or at low temperature (−4°C) from our previous study (NCBI BioProject accession: PRJNA685002) were reanalyzed [46]. Briefly, fastp was used for quality filtering, FastQC v0.11.7 [47] was used for quality control, and Bismark (v0.23.0) [48] was used to align the trimmed reads to the assembled NTS57 reference genome. The DNA methylation levels for individual CpGs, CHG, and CHH sites (H = A, T, or C) were calculated separately using methyKit (v1.22.0) [49]; only cytosine sites with ≥10 covered sequencing reads were retained. Methylated reads (containing Cs) and unmethylated reads (containing Ts) at each cytosine site were counted, and the percentage of methylated reads among the total reads covering the corresponding cytosine was calculated to quantify the DNA methylome for each sample at single base resolution. To identify differentially methylated regions, we used a window-based method to calculate the overall methylation level between samples for comparisons. For this purpose, 1,000-bp continuous windows were generated across 19 chromosomes of the NTS57 genome. Windows that covered ≥ 10 reads that showed ≥ 25% methylation level differences with q-value ≤ 0.01 were considered as differentially methylated regions using methyKit (v1.22.0) [49].

To compare the overall methylation levels of upstream and downstream regions, gene body of protein-coding genes, and repeat elements, the mean CpG, CHG, and CHH methylation levels for the 2-kb upstream and downstream regions of each type of genomic feature were binned into 100 equal-length windows, and the mean methylation levels in each window were calculated using a custom R script.

#### Structural variation (SV) identification

For SV calling, minimap2 (v2.21-r1071) [50] was used to generate pairwise genome alignments between the genomes of each of the nine rapeseed accessions and the NTS57 genome using parameters -ax asm5 -eqx. The alignment results were passed to SyRI [51] and SVs consisting of insertions, deletions, inversions (< 1 Mb in size), translocations (> 50 bp in length), or duplications were retained. SVs containing N sequences and ambitious alignment margins and/or poor synteny alignment around the breakpoint were filtered. SVs between different genomes were visualized using plotsr [52].

### Supplementary figures

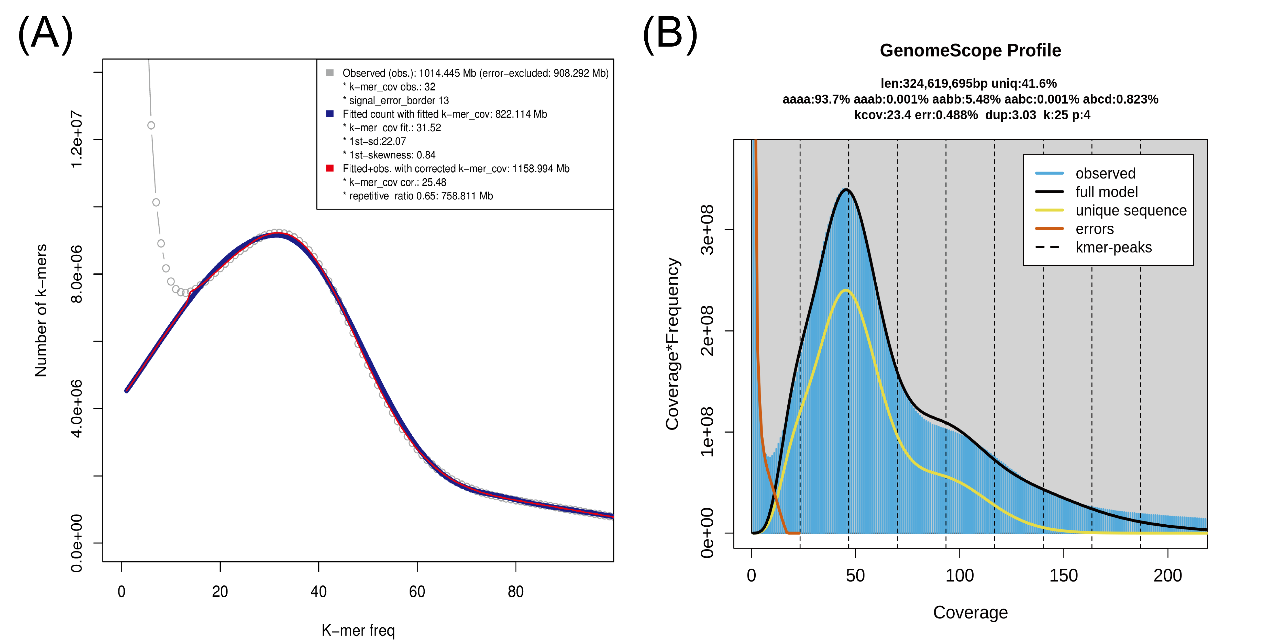

**Figure S1. Genome size estimation based on survey sequencing.** (A) K-mer frequency histogram produced by findGSE using a 25-mer k-mer. Gray line, observed k-mer frequency; blue line, fitted model without k-mer correction; red line, fitted model with k-mer correction. The fitted model was used to estimate the genome size. (B) GenomeScope2 plot of the 25-mer k-mer frequency for sequencing reads of NTS57.

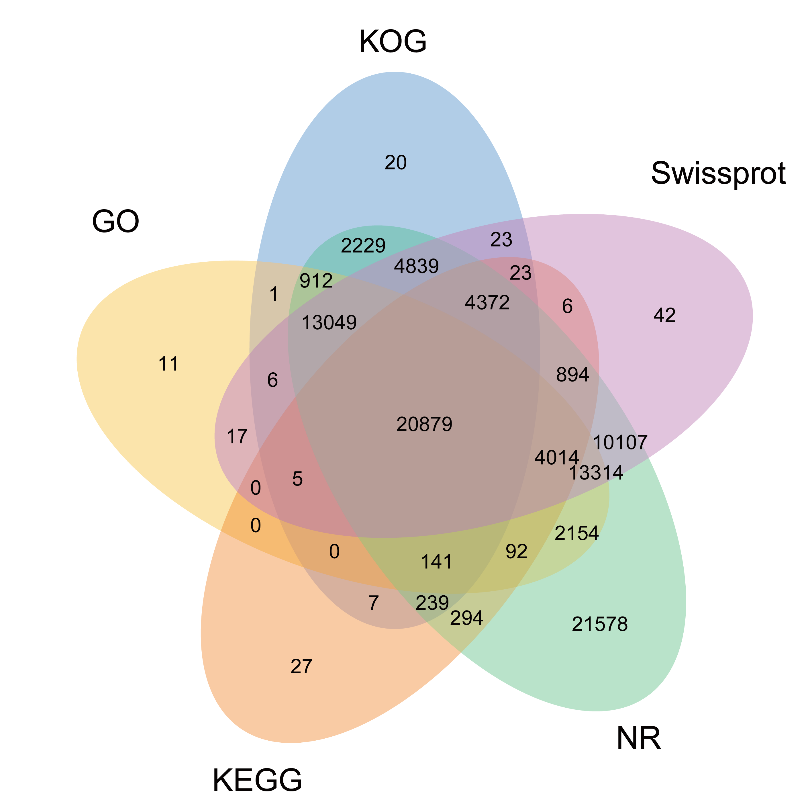

**Figure S2. Venn diagram of gene function annotations obtained using several protein-related databases.** NR, NCBI nonredundant protein database; KEGG, Kyoto Encyclopedia of Genes and Genomes; KOG, Eukaryotic Orthologous Groups; GO, gene ontology.

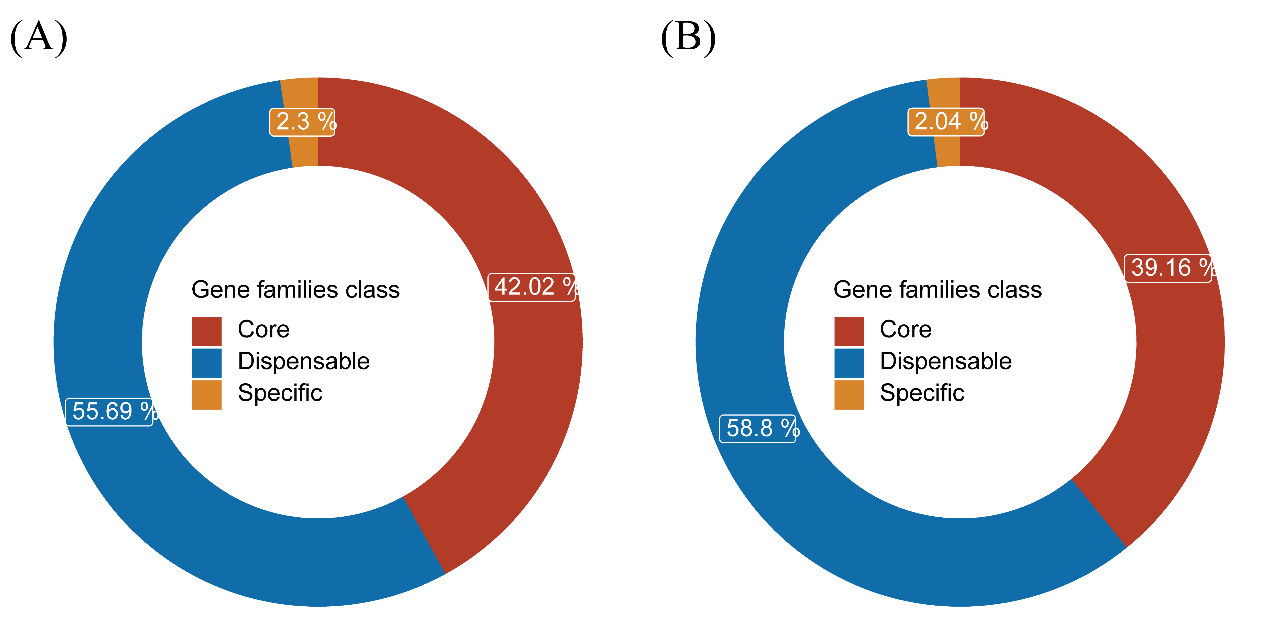
**Figure S3. Percentage of core genes, dispensable genes, and accession-specific genes in the rapeseed pangenome**. (A) and (B) show the percentage of three classes of gene families in the A and C subgenomes of rapeseed, respectively.

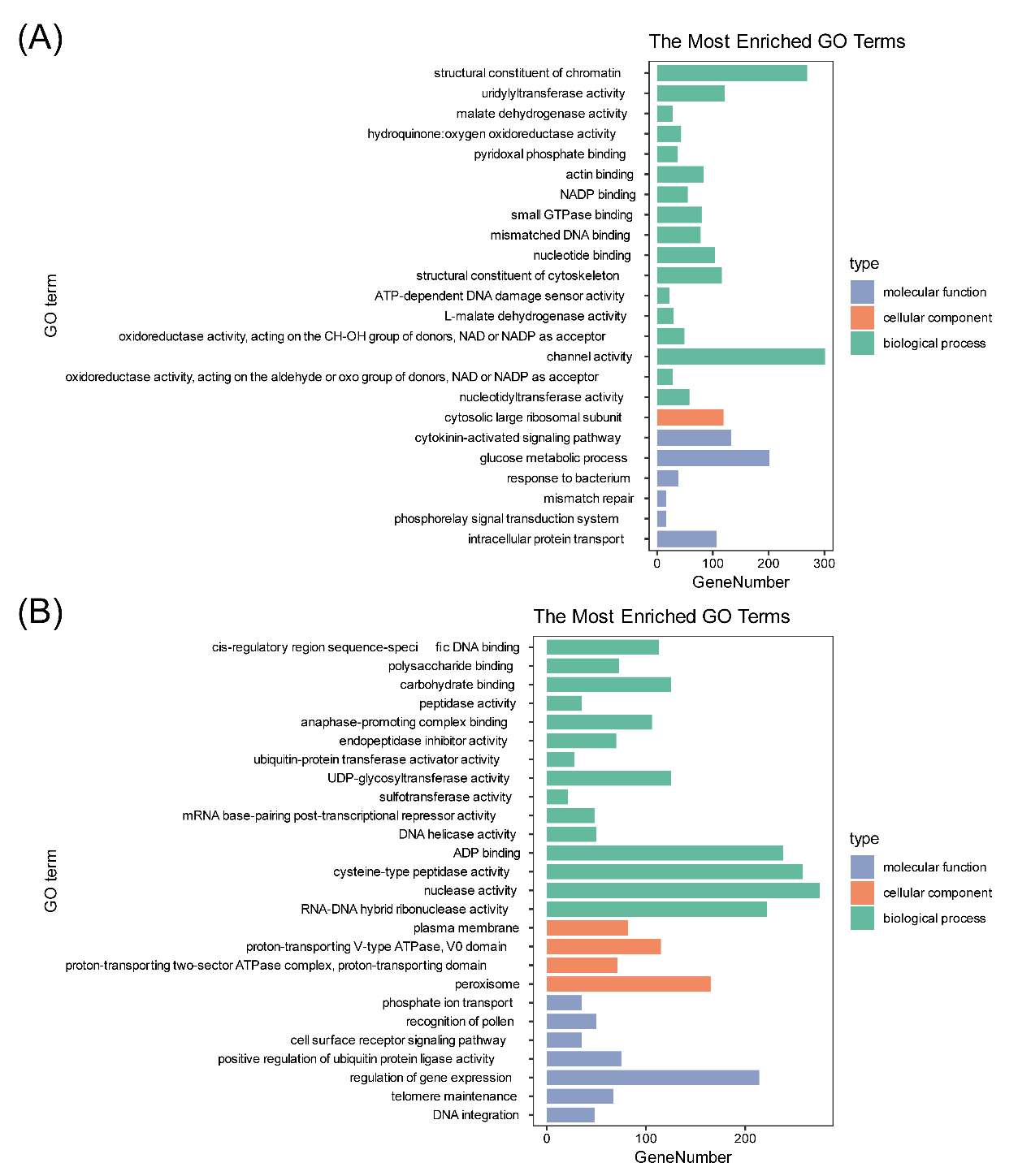

**Figure S4. Gene function enrichment for core and dispensable genes of NTS57.** (A) Gene function enrichment for core genes shared by all rapeseed accessions examined (B) Gene function enrichment for dispensable genes; i.e., genes that are not shared by all rapeseed accessions.

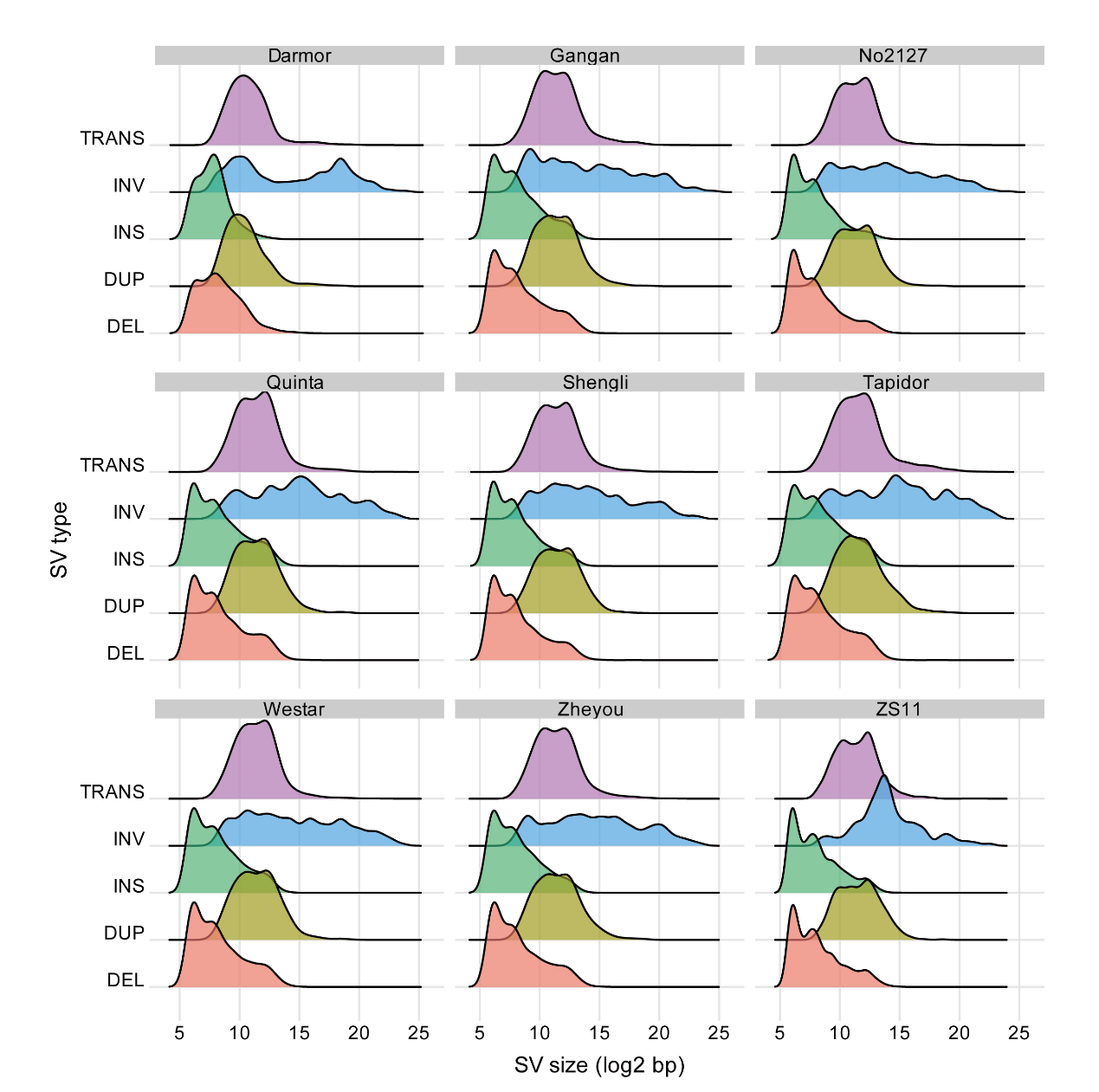

**Figure S5. Length statistics for structural variations (SVs) in different rapeseed accessions using NTS57 as the reference genome.** TRANS, INV, INS, DUP, and DEL, translocation, inversion, insertion, duplication, and deletion, respectively.

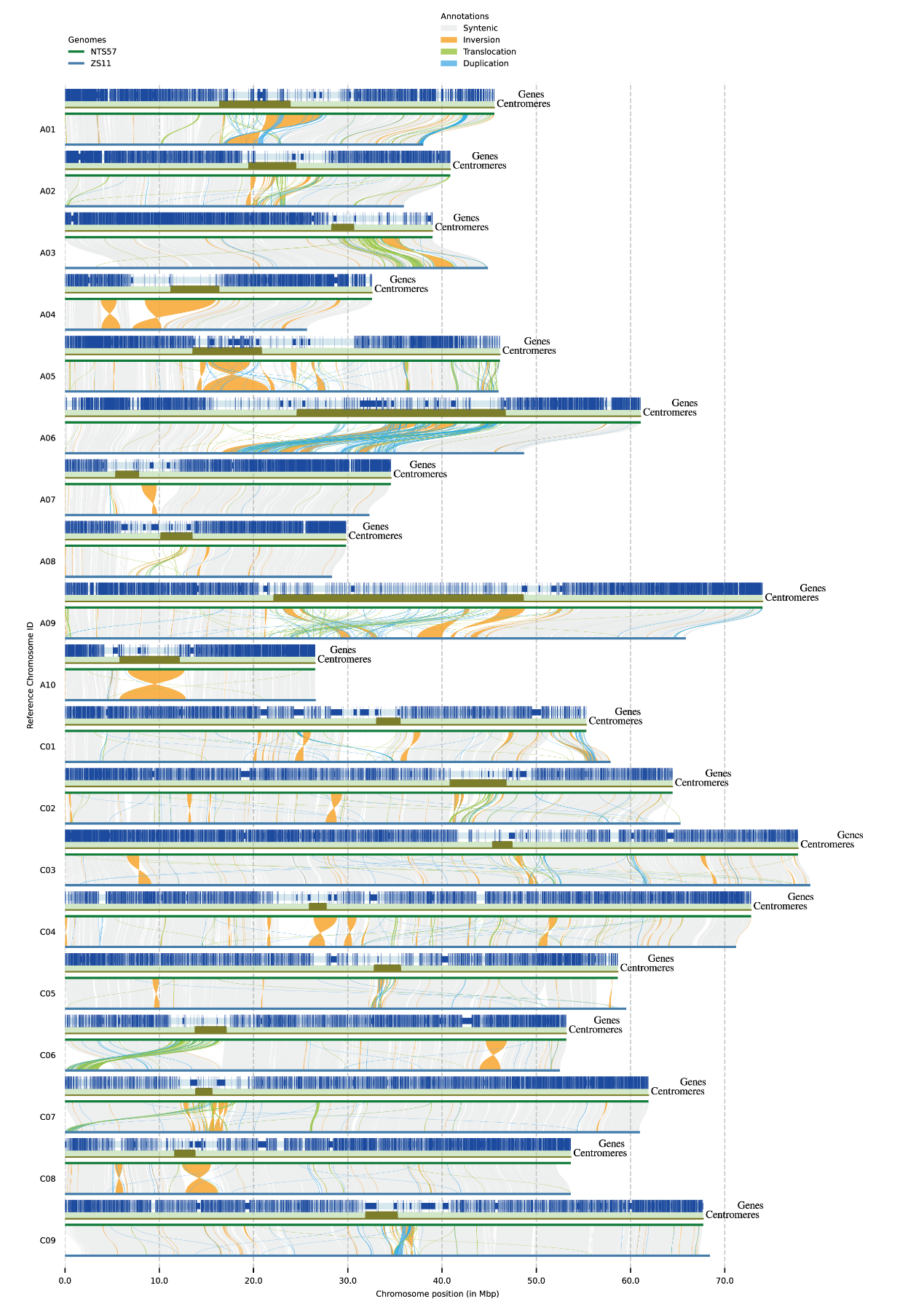
**Figure S6. Synteny and rearrangement plot for 19 chromosomes of NTS57 and ZS11.** The plot was obtained using SyRi. Positions of protein-coding genes and centromeres are shown in blue and dark green genomic tracks, respectively.

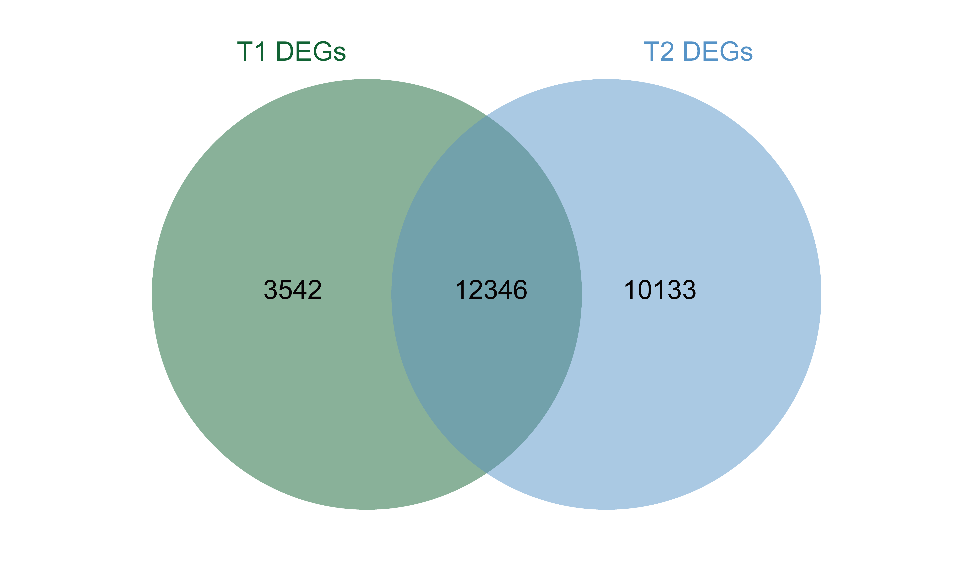

**Figure S7. Number of DEGs identified in NTS57 under cold stress.** "T1" and "T2" represent the samples collected 6 and 24 hours after cold stress, respectively.

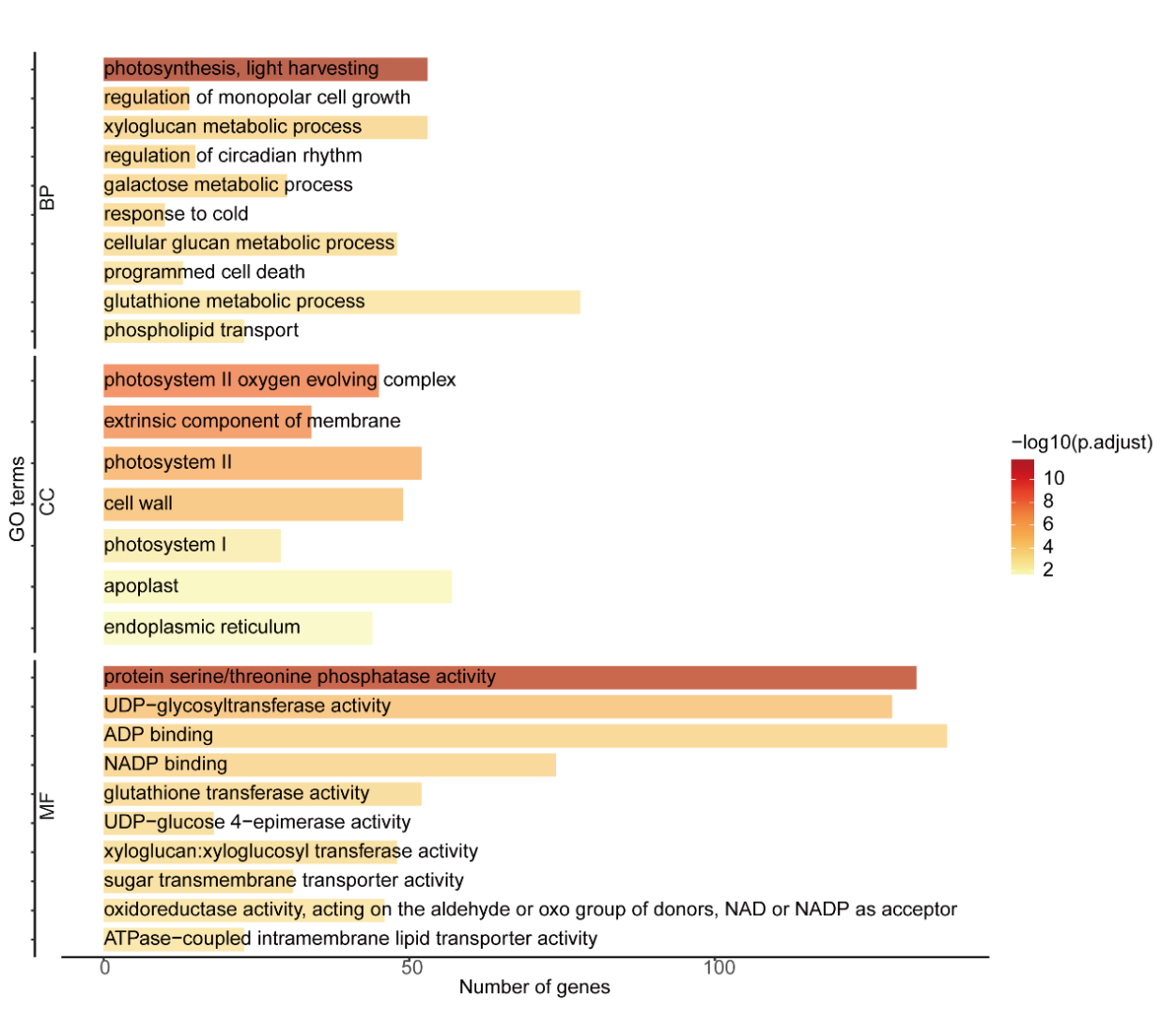

**Figure S8.** **Gene ontology (GO) functional enrichment for the DEGs identified in T2 samples relative to those in the normal condition samples.** T2, leaf samples taken after 24 h at −4°C. BP, biological process; CC, cellular component; MF, molecular function.

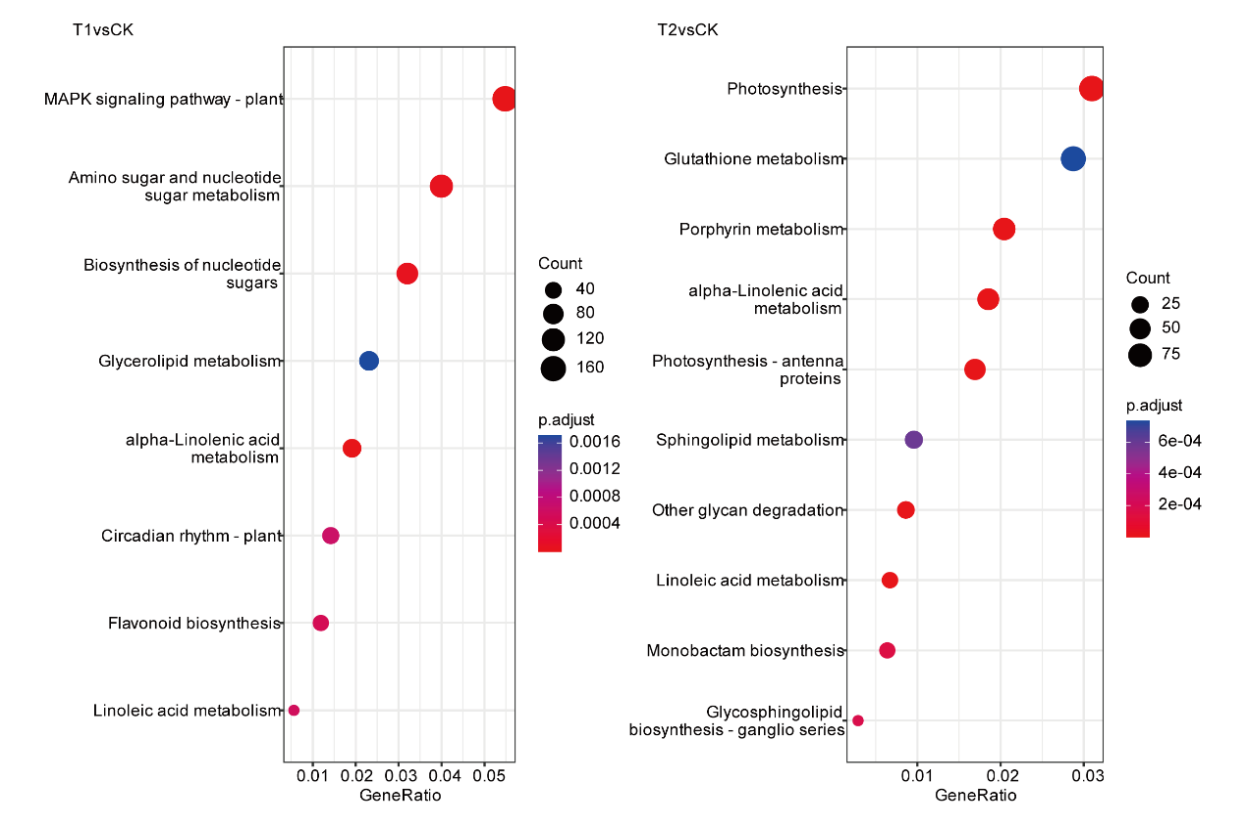

**Figure S9. KEGG pathway enrichment for the DEGs identified in T1 or T2 samples relative to** **those in normal condition (CK) samples.** T1 and T2, leaf samples taken after 12 h and 24 h at −4°C, respectively.

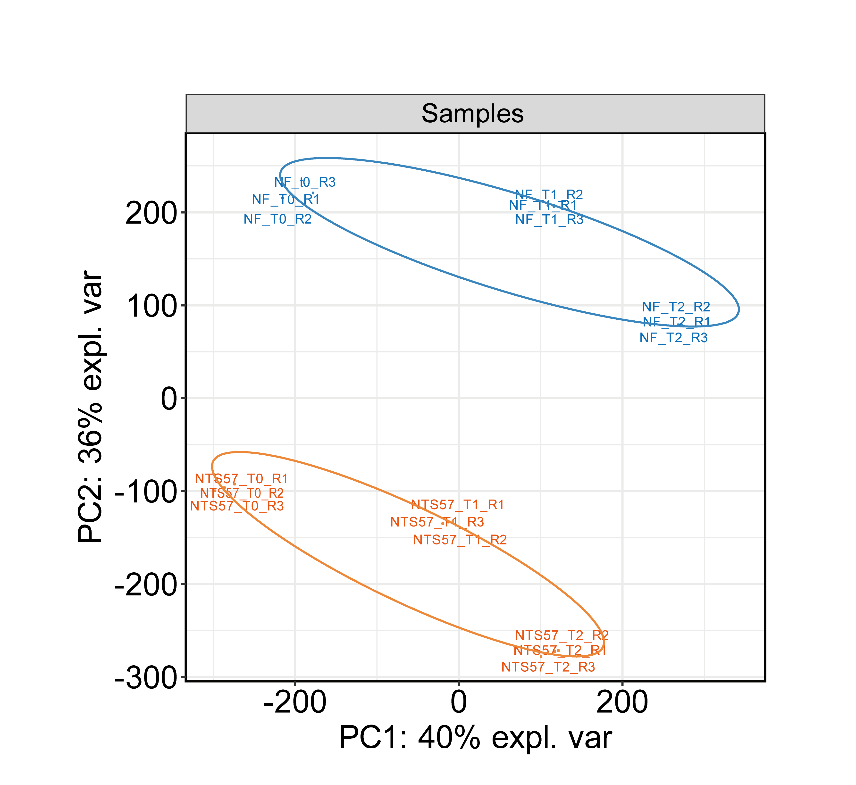

**Figure S10. Principal component analysis of gene expression profiles from accessions NTS57 and NF under cold stress.** PC1 and PC2, principal components 1 and 2.

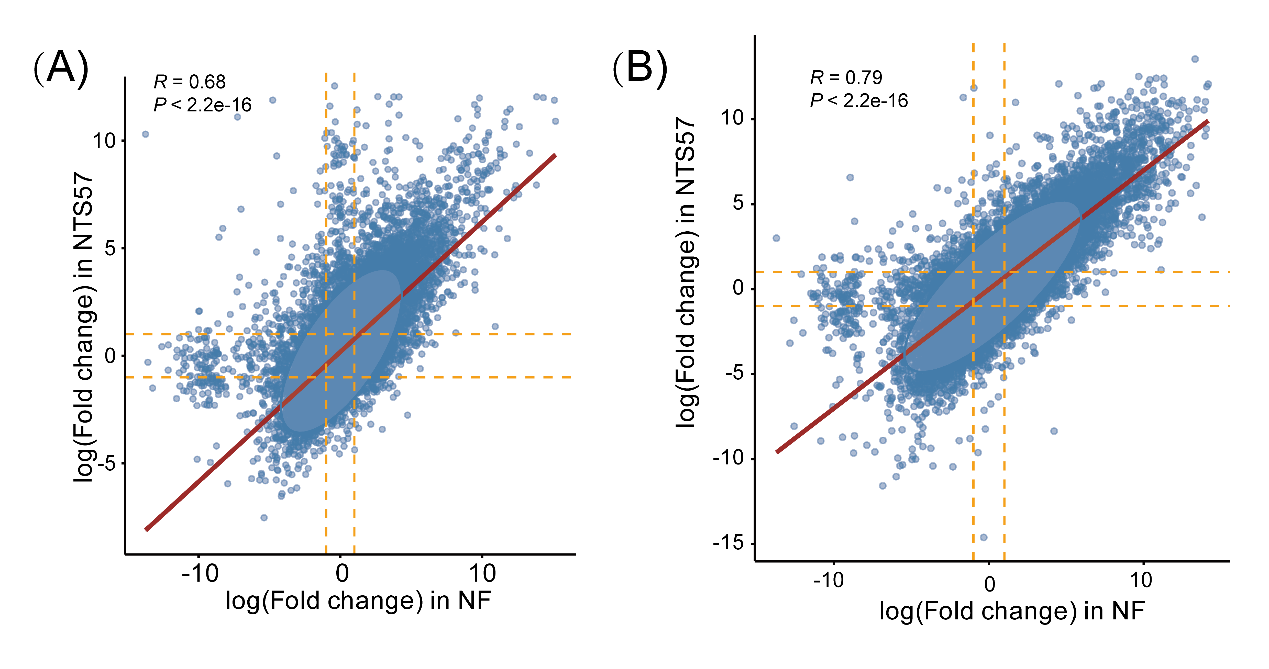
**Figure S11. Gene expression changes in NTS57 and NF under cold stress.** (A) and (B) show the gene expression changes of two accessions in T1 and T2 samples, respectively.

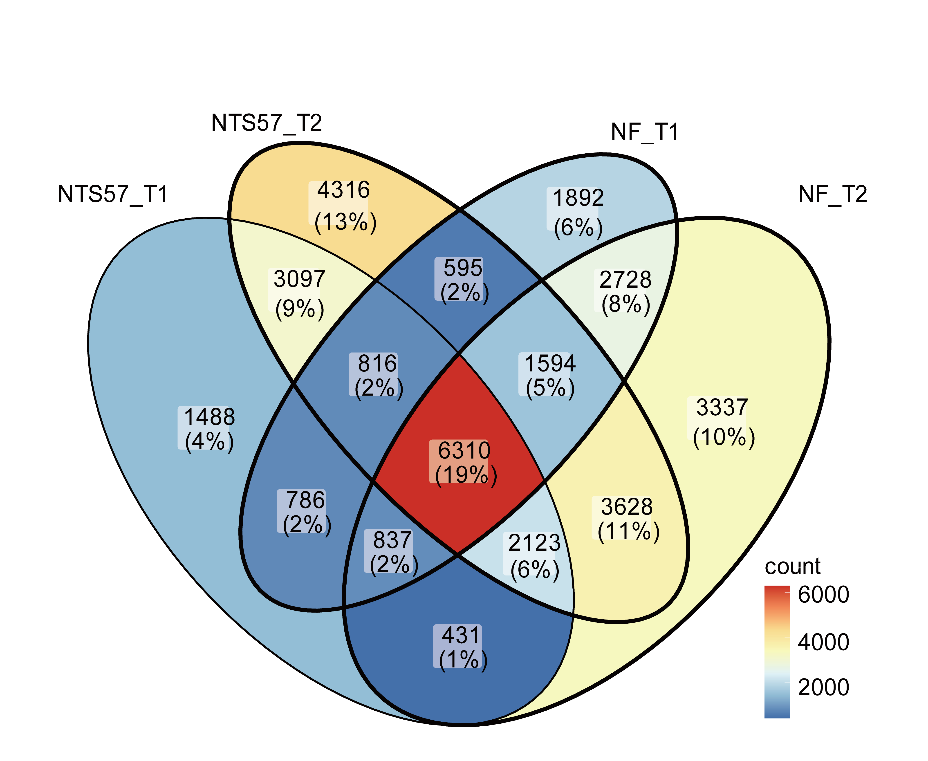

**Figure S12. Veen diagram showing the intersection of the DEGs in NTS57 and NF in the T1 and T2 samples.** "T1" and "T2" represent the samples collected 6 and 24 hours after cold stress, respectively.

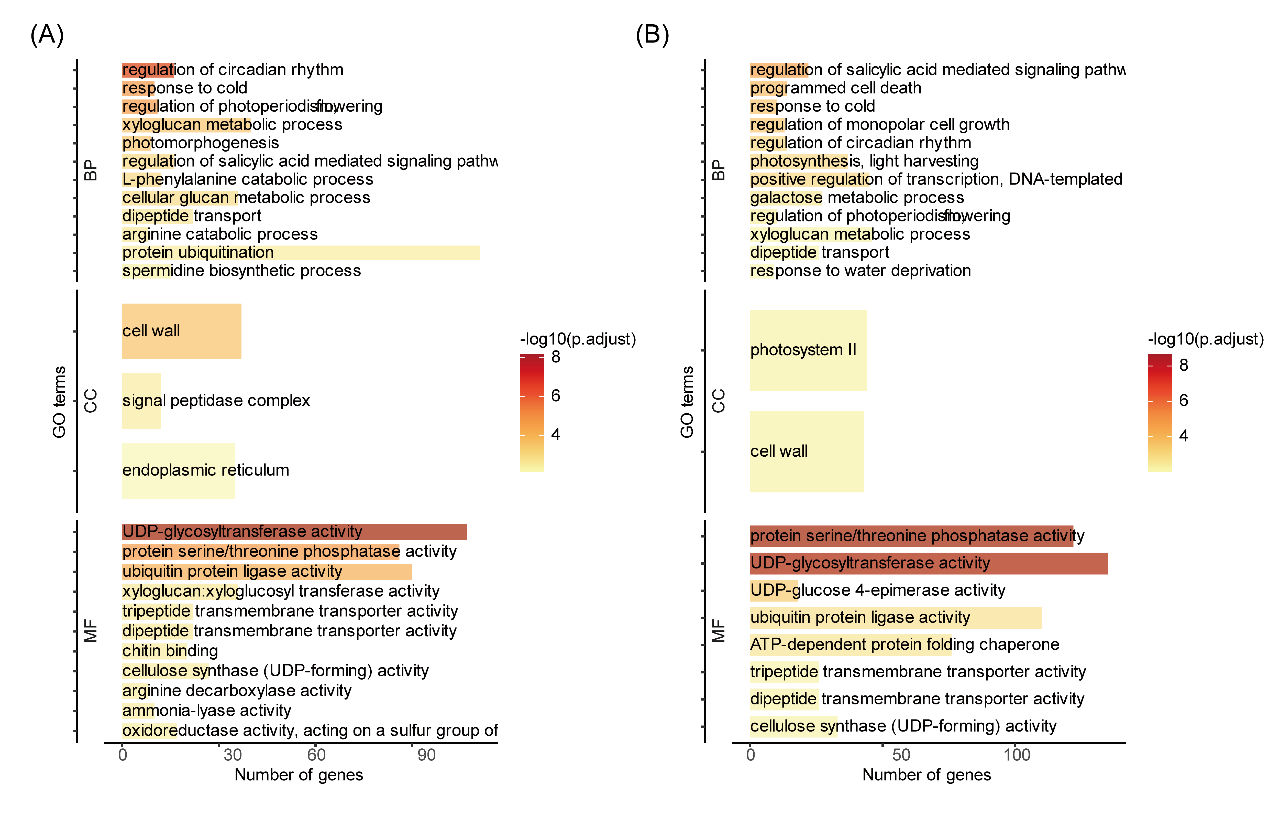
**Figure S13. Gene functional enrichment of the DEGs in NF in the T1 (A) and T2 (B) samples.** BP, biological process; CC, cellular component; MF, molecular function.

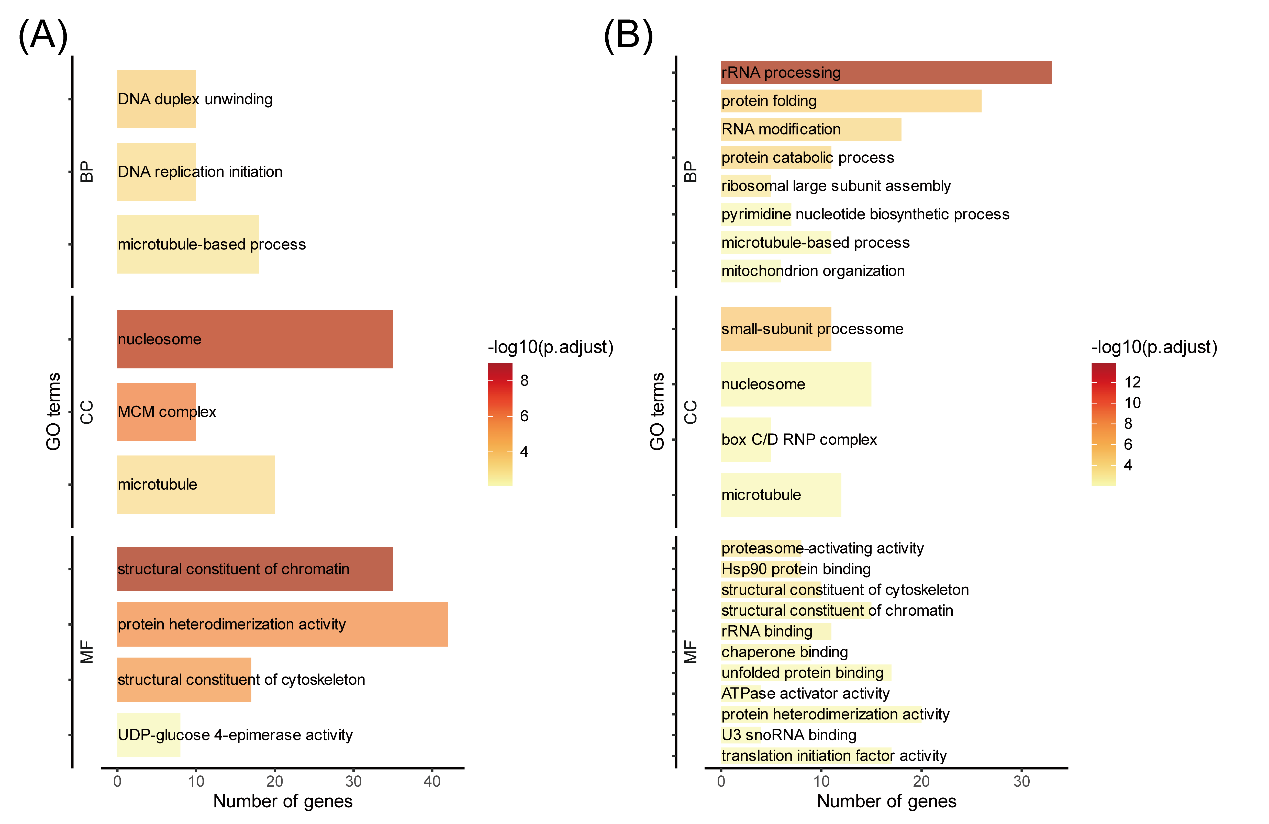

**Figure S14. Gene functional enrichment of the** **NTS57-specific differentially expressed genes (DEGs) of NTS57 in the T1 (A) and T2 (B) samples.** BP, biological process; CC, cellular component; MF, molecular function.

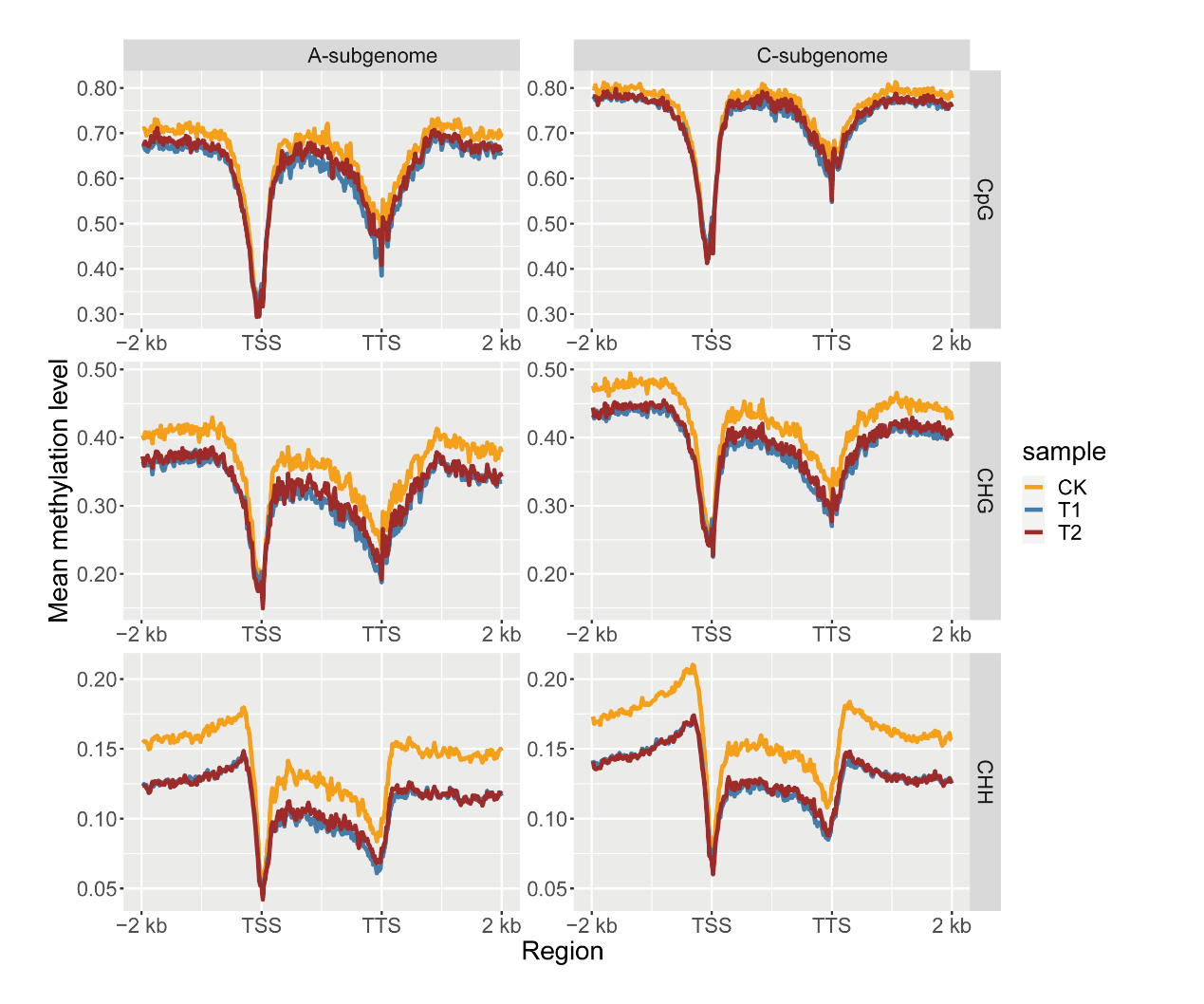

**Figure S15. Methylation level profiles of the A and C subgenomes of NTS57 under cold stress (T1 and T2) and normal conditions (CK).** The gene body was divided into 100 bins of equal size. For each bin in each gene, the methylation level was calculated by dividing the total number of methylated Cs by the total number of Cs, and the average of methylation level in the window across all protein-coding genes was used to generate the profiles. The three methylation profiles for CpG, CHG, and CHH contexts were analyzed independently. TSS, transcription start site; TTS, transcription termination site.

**REFERENCES**

1. Kokot, Marek, Maciej Dlugosz, Sebastian Deorowicz. 2017. “KMC 3: counting and manipulating k-mer statistics.” *Bioinformatics* 33: 2759-2761. <https://doi.org/10.1093/bioinformatics/btx304>

2. Sun, Hequan, Jia Ding, Mathieu Piednoël, Korbinian Schneeberger. 2018. “findGSE: estimating genome size variation within human and Arabidopsis using k-mer frequencies.” *Bioinformatics* 34: 550-557. <https://doi.org/10.1093/bioinformatics/btx637>

3. Ranallo-Benavidez, T Rhyker, Kamil S Jaron, Michael C Schatz. 2020. “GenomeScope 2.0 and Smudgeplot for reference-free profiling of polyploid genomes.” *Nat Commun* 11: 1432. <https://doi.org/10.1038/s41467-020-14998-3>

4. Cheng, Haoyu, Gregory T Concepcion, Xiaowen Feng, Haowen Zhang, Heng Li. 2021. “Haplotype-resolved de novo assembly using phased assembly graphs with hifiasm.” *Nat Methods* 18: 170-175. <https://doi.org/10.1038/s41592-020-01056-5>

5. Hu, Jiang, Junpeng Fan, Zongyi Sun, Shanlin Liu. 2020. “NextPolish: a fast and efficient genome polishing tool for long-read assembly.” *Bioinformatics* 36: 2253-2255. <https://doi.org/10.1093/bioinformatics/btz891>

6. Huang, Shengfeng, Mingjing Kang, Anlong Xu. 2017. “HaploMerger2: rebuilding both haploid sub-assemblies from high-heterozygosity diploid genome assembly.” *Bioinformatics* 33: 2577-2579. <https://doi.org/10.1093/bioinformatics/btx220>

7. Chen, Shifu, Yanqing Zhou, Yaru Chen, Jia Gu. 2018. “fastp: an ultra-fast all-in-one FASTQ preprocessor.” *Bioinformatics* 34: i884-i890. <https://doi.org/10.1093/bioinformatics/bty560>

8. Langmead, Ben, Steven L Salzberg. 2012. “Fast gapped-read alignment with Bowtie 2.” *Nat Methods* 9: 357-359. <https://doi.org/10.1038/nmeth.1923>

9. Servant, Nicolas, Nelle Varoquaux, Bryan R Lajoie, Eric Viara, Chong-Jian Chen, Jean-Philippe Vert, Edith Heard, Job Dekker, Emmanuel Barillot. 2015. “HiC-Pro: an optimized and flexible pipeline for Hi-C data processing.” *Genome Biol* 16: 259. <https://doi.org/10.1186/s13059-015-0831-x>

10. Burton, Joshua N, Andrew Adey, Rupali P Patwardhan, Ruolan Qiu, Jacob O Kitzman, Jay Shendure. 2013. “Chromosome-scale scaffolding of de novo genome assemblies based on chromatin interactions.” *Nat Biotechnol* 31: 1119-1125. <https://doi.org/10.1038/nbt.2727>

11. Simao, Felipe A, Robert M Waterhouse, Panagiotis Ioannidis, Evgenia V Kriventseva, Evgeny M Zdobnov. 2015. “BUSCO: assessing genome assembly and annotation completeness with single-copy orthologs.” *Bioinformatics* 31: 3210-3212. <https://doi.org/10.1093/bioinformatics/btv351>

12. Li, Heng, Richard Durbin. 2009. “Fast and accurate short read alignment with Burrows-Wheeler transform.” *Bioinformatics* 25: 1754-1760. <https://doi.org/10.1093/bioinformatics/btp324>

13. Li, Heng, Bob Handsaker, Alec Wysoker, Tim Fennell, Jue Ruan, Nils Homer, Gabor Marth, Goncalo Abecasis, Richard Durbin. 2009. “The Sequence Alignment/Map format and SAMtools.” *Bioinformatics* 25: 2078-2079. <https://doi.org/10.1093/bioinformatics/btp352>

14. Danecek, Petr, Shane A McCarthy. 2017. “BCFtools/csq: haplotype-aware variant consequences.” *Bioinformatics* 33: 2037-2039. <https://doi.org/10.1093/bioinformatics/btx100>

15. Kim, Daehwan, Joseph M Paggi, Chanhee Park, Christopher Bennett, Steven L Salzberg. 2019. “Graph-based genome alignment and genotyping with HISAT2 and HISAT-genotype.” *Nat Biotechnol* 37: 907-915. <https://doi.org/10.1038/s41587-019-0201-4>

16. Wang, Xuewen, Le Wang. 2016. “GMATA: An Integrated Software Package for Genome-Scale SSR Mining, Marker Development and Viewing.” *Front Plant Sci* 7: 1350. <https://doi.org/10.3389/fpls.2016.01350>

17. Benson, Gary. 1999. “Tandem repeats finder: a program to analyze DNA sequences.” *Nucleic Acids Res* 27: 573-580. <https://doi.org/10.1093/nar/27.2.573>

18. Han, Yujun, Susan R Wessler. 2010. “MITE-Hunter: a program for discovering miniature inverted-repeat transposable elements from genomic sequences.” *Nucleic Acids Res* 38: e199. <https://doi.org/10.1093/nar/gkq862>

19. Flynn, Jullien M, Robert Hubley, Clement Goubert, Jeb Rosen, Andrew G Clark, Cedric Feschotte, Arian F Smit. 2020. “RepeatModeler2 for automated genomic discovery of transposable element families.” *Proc Natl Acad Sci U S A* 117: 9451-9457. <https://doi.org/10.1073/pnas.1921046117>

20. Ou, Shujun, Ning Jiang. 2018. “LTR_retriever: A Highly Accurate and Sensitive Program for Identification of Long Terminal Repeat Retrotransposons.” *Plant Physiol* 176: 1410-1422. <https://doi.org/10.1104/pp.17.01310>

21. Tarailo-Graovac, Maja, Nansheng Chen. 2009. “Using RepeatMasker to identify repetitive elements in genomic sequences.” *Curr Protoc Bioinformatics* Chapter 4: 4 10 11-14 10 14. <https://doi.org/10.1002/0471250953.bi0410s25>

22. Lin, Yunzhi, Chen Ye, Xingzhu Li, Qinyao Chen, Ying Wu, Feng Zhang, Rui Pan, et al. 2023. “quarTeT: a telomere-to-telomere toolkit for gap-free genome assembly and centromeric repeat identification.” *Hortic Res* 10: uhad127. <https://doi.org/10.1093/hr/uhad127>

23. Keilwagen, Jens, Frank Hartung, Jan Grau. 2019. “GeMoMa: Homology-Based Gene Prediction Utilizing Intron Position Conservation and RNA-seq Data.” *Methods Mol Biol* 1962: 161-177. <https://doi.org/10.1007/978-1-4939-9173-0_9>

24. Dobin, Alexander, Carrie A Davis, Felix Schlesinger, Jorg Drenkow, Chris Zaleski, Sonali Jha, Philippe Batut, Mark Chaisson, Thomas R Gingeras. 2013. “STAR: ultrafast universal RNA-seq aligner.” *Bioinformatics* 29: 15-21. <https://doi.org/10.1093/bioinformatics/bts635>

25. Pertea, Mihaela, Geo M Pertea, Corina M Antonescu, Tsung-Cheng Chang, Joshua T Mendell, Steven L Salzberg. 2015. “StringTie enables improved reconstruction of a transcriptome from RNA-seq reads.” *Nat Biotechnol* 33: 290-295. <https://doi.org/10.1038/nbt.3122>

26. Stanke, Mario, Rasmus Steinkamp, Stephan Waack, Burkhard Morgenstern. 2004. “AUGUSTUS: a web server for gene finding in eukaryotes.” *Nucleic Acids Res* 32: W309-312. <https://doi.org/10.1093/nar/gkh379>

27. Haas, Brian J, Steven L Salzberg, Wei Zhu, Mihaela Pertea, Jonathan E Allen, Joshua Orvis, Owen White, C Robin Buell, Jennifer R Wortman. 2008. “Automated eukaryotic gene structure annotation using EVidenceModeler and the Program to Assemble Spliced Alignments.” *Genome Biol* 9: R7. <https://doi.org/10.1186/gb-2008-9-1-r7>

28. Boutet, Emmanuel, Damien Lieberherr, Michael Tognolli, Michel Schneider, Amos Bairoch. 2007. “UniProtKB/Swiss-Prot.” *Methods Mol Biol* 406: 89-112. <https://doi.org/10.1007/978-1-59745-535-0_4>

29. Pruitt, Kim D, Tatiana Tatusova, Donna R Maglott. 2007. “NCBI reference sequences (RefSeq): a curated non-redundant sequence database of genomes, transcripts and proteins.” *Nucleic Acids Res* 35: D61-65. <https://doi.org/10.1093/nar/gkl842>

30. Kanehisa, Minoru, Susumu Goto. 2000. “KEGG: kyoto encyclopedia of genes and genomes.” *Nucleic Acids Res* 28: 27-30. <https://doi.org/10.1093/nar/28.1.27>

31. Tatusov, Roman L., Darren A. Natale, Igor V. Garkavtsev, Tatiana A. Tatusova, Uma T. Shankavaram, Bachoti S. Rao, Boris Kiryutin, Michael Y. Galperin, Natalie D. Fedorova, Eugene V. Koonin. 2001. “The COG database: new developments in phylogenetic classification of proteins from complete genomes.” *Nucleic Acids Res* 29: 22-28. <https://doi.org/10.1093/nar/29.1.22>

32. Gene Ontology, Consortium. 2015. “Gene Ontology Consortium: going forward.” *Nucleic Acids Res* 43: D1049-1056. <https://doi.org/10.1093/nar/gku1179>

33. Quevillon, E., V. Silventoinen, S. Pillai, N. Harte, N. Mulder, R. Apweiler, R. Lopez. 2005. “InterProScan: protein domains identifier.” *Nucleic Acids Res* 33: W116-120. <https://doi.org/10.1093/nar/gki442>

34. Altschul, Stephen F., Warren Gish, Webb Miller, Eugene W. Myers, David J. Lipman. 1990. “Basic local alignment search tool.” *J Mol Biol* 215: 403-410. <https://doi.org/10.1016/S0022-2836(05)80360-2>

35. Chan, Patricia P, Brian Y Lin, Allysia J Mak, Todd M Lowe. 2021. “tRNAscan-SE 2.0: improved detection and functional classification of transfer RNA genes.” *Nucleic Acids Res* 49: 9077-9096. <https://doi.org/10.1093/nar/gkab688>

36. Nawrocki, Eric P, Sean R Eddy. 2013. “Infernal 1.1: 100-fold faster RNA homology searches.” *Bioinformatics* 29: 2933-2935. <https://doi.org/10.1093/bioinformatics/btt509>

37. Lagesen, Karin, Peter Hallin, Einar Andreas Rodland, Hans-Henrik Staerfeldt, Torbjorn Rognes, David W Ussery. 2007. “RNAmmer: consistent and rapid annotation of ribosomal RNA genes.” *Nucleic Acids Res* 35: 3100-3108. <https://doi.org/10.1093/nar/gkm160>

38. Tang, Haibao, John E Bowers, Xiyin Wang, Ray Ming, Maqsudul Alam, Andrew H Paterson. 2008. “Synteny and collinearity in plant genomes.” *Science* 320: 486-488. <https://doi.org/10.1126/science.1153917>

39. Emms, David M, Steven Kelly. 2019. “OrthoFinder: phylogenetic orthology inference for comparative genomics.” *Genome Biol* 20: 238. <https://doi.org/10.1186/s13059-019-1832-y>

40. Katoh, Kazutaka, Daron M Standley. 2013. “MAFFT multiple sequence alignment software version 7: improvements in performance and usability.” *Mol Biol Evol* 30: 772-780. <https://doi.org/10.1093/molbev/mst010>

41. Nguyen, Lam-Tung, Heiko A Schmidt, Arndt von Haeseler, Bui Quang Minh. 2015. “IQ-TREE: a fast and effective stochastic algorithm for estimating maximum-likelihood phylogenies.” *Mol Biol Evol* 32: 268-274. <https://doi.org/10.1093/molbev/msu300>

42. Kalyaanamoorthy, Subha, Bui Quang Minh, Thomas K F Wong, Arndt von Haeseler, Lars S Jermiin. 2017. “ModelFinder: fast model selection for accurate phylogenetic estimates.” *Nat Methods* 14: 587-589. <https://doi.org/10.1038/nmeth.4285>

43. Bolger, Anthony M, Marc Lohse, Bjoern Usadel. 2014. “Trimmomatic: a flexible trimmer for Illumina sequence data.” *Bioinformatics* 30: 2114-2120. <https://doi.org/10.1093/bioinformatics/btu170>

44. Wei, Jiaping, Guoqiang Zheng, Xingwang Yu, Sushuang Liu, Xiaoyun Dong, Xiaodong Cao, Xinling Fang, et al. 2021. “Comparative Transcriptomics and Proteomics Analyses of Leaves Reveals a Freezing Stress-Responsive Molecular Network in Winter Rapeseed (*Brassica rapa* L.).” *Front Plant Sci* 12: 664311. <https://doi.org/10.3389/fpls.2021.664311>

45. Anders, Simon, Davis J McCarthy, Yunshun Chen, Michal Okoniewski, Gordon K Smyth, Wolfgang Huber, Mark D Robinson. 2013. “Count-based differential expression analysis of RNA sequencing data using R and Bioconductor.” *Nature Protocols* 8: 1765-1786. <https://doi.org/10.1038/nprot.2013.099>

46. Wei, Jiaping, Yingzi Shen, Xiaoyun Dong, Yajing Zhu, Junmei Cui, Hui Li, Guoqiang Zheng, Haiyan Tian, Ying Wang, Zigang Liu. 2022. “DNA methylation affects freezing tolerance in winter rapeseed by mediating the expression of genes related to JA and CK pathways.” *Front Genet* 13: 968494. <https://doi.org/10.3389/fgene.2022.968494>

47. Wingett, Steven W, Simon Andrews. 2018. “FastQ Screen: A tool for multi-genome mapping and quality control.” *F1000Res* 7: 1338. <https://doi.org/10.12688/f1000research.15931.2>

48. Krueger, Felix, Simon R Andrews. 2011. “Bismark: a flexible aligner and methylation caller for Bisulfite-Seq applications.” *Bioinformatics* 27: 1571-1572. <https://doi.org/10.1093/bioinformatics/btr167>

49. Akalin, Altuna, Matthias Kormaksson, Sheng Li, Francine E Garrett-Bakelman, Maria E Figueroa, Ari Melnick, Christopher E Mason. 2012. “methylKit: a comprehensive R package for the analysis of genome-wide DNA methylation profiles.” *Genome Biol* 13: R87. <https://doi.org/10.1186/gb-2012-13-10-r87>

50. Li, Heng. 2018. “Minimap2: pairwise alignment for nucleotide sequences.” *Bioinformatics* 34: 3094-3100. <https://doi.org/10.1093/bioinformatics/bty191>

51. Goel, Manish, Hequan Sun, Wen-Biao Jiao, Korbinian Schneeberger. 2019. “SyRI: finding genomic rearrangements and local sequence differences from whole-genome assemblies.” *Genome Biol* 20: 277. <https://doi.org/10.1186/s13059-019-1911-0>

52. Goel, Manish, Korbinian Schneeberger. 2022. “plotsr: visualizing structural similarities and rearrangements between multiple genomes.” *Bioinformatics* 38: 2922-2926. <https://doi.org/10.1093/bioinformatics/btac196>
